# Adiponectin-receptor agonism prevents right ventricular tissue pathology in a mouse model of Duchenne muscular dystrophy

**DOI:** 10.1101/2024.10.23.619868

**Authors:** Shivam Gandhi, Luca J. Delfinis, Parashar D. Bhatt, Madison C. Garibotti, Catherine A. Bellissimo, Amireza N. Goli, Brooke A. Morris, Aditya N. Brahmbhatt, Simona Yakobov-Shimonov, Fasih A. Rahman, Joe Quadrilatero, Jeremy A. Simpson, Gary Sweeney, Ali A. Abdul-Sater, Peter H. Backx, Henry H. Hsu, Christopher G.R. Perry

## Abstract

Cardiac fibrosis during Duchenne muscular dystrophy (DMD) arises from cellular damage and inflammation and is associated with myofibre hypertrophy and metabolic dysfunction. However, the extent to which these relationships develop across all 4 cardiac chambers, particularly during early-stage disease, remains unknown. Here, we discovered that very young D2.*mdx* mice exhibit fibrosis exclusively in the right ventricle (RV) and left atrium. Concurrent cardiomyocyte hypertrophy and disorganization in the RV were related to a highly specific inflammatory signature of increased infiltrating pro-inflammatory macrophages (CD11b^+^CD45^+^CD64^+^F4/80^+^CCR2^+^), myofibre mitochondrial-linked apoptosis, and reduced carbohydrate and fat oxidation. This relationship did not occur in the left ventricle. Short-term daily administration of a peptidomimetic adiponectin receptor agonist, ALY688, completely prevented RV fibrosis, myofibre hypertrophy, infiltrating macrophages and mitochondrial stress as well as left atrial fibrosis. Our discoveries demonstrate early-stage cardiac tissue pathology occurs in a chamber-specific manner and is prevented by adiponectin receptor agonism, thereby opening a new direction for developing therapies that prevent tissue remodeling during DMD.

**Teaser:** Heterogeneous distribution of fibrosis, inflammation and metabolic dysfunction across the heart is prevented by adiponectin receptor agonism in a mouse model of Duchenne muscular dystrophy.

## Introduction

Duchenne muscular dystrophy (DMD) is a devasting X-linked recessive neuromuscular disease with an approximate global birth prevalence of 19.8 per 100,000 live male births (1). It is caused by mutations to the *DMD* gene, which encodes the subsarcolemmal protein dystrophin (2–4). Absence of dystrophin in cardiomyocytes can hinder the cardiac extracellular matrix (ECM) from anchoring to the actin and microtubule (5) cytoskeleton networks – a process critical in providing stability to the sarcolemmal membrane during contraction-relaxation cycles. This can lead to contraction-induced membrane damage that leads to repeated cycles of systemic inflammation, necrosis, and fibrosis as well as eventual heart failure (6).

Cardiac fibrosis is a programmed event triggered by repeated injury or damage. While fibrosis contributes to supporting the structural integrity of the heart by replacing dead or dying tissue with collagenous scar tissue (7–9), it also interferes with heart function and is linked to cardiomyocyte disorganization, hypertrophy, and inflammation (7,9–11). As such, there is extensive interest in developing anti-fibrotics as a means of preventing pathological tissue remodeling in DMD. However, therapy development would be better informed by understanding how fibrosis develops in each chamber of the heart, particularly during the early stages of disease. While many clinical studies have traditionally focused on LV fibrosis (9,12–15) given this chamber demonstrates functional decrements in later stages of disease (16–18), some post-mortem case reports have also identified right ventricular fibrosis (12,19). The potential for atrial remodeling is even less understood with at least one case study showing fibrosis in the left atrium that was not seen in the right atrium (20). While these observations suggest that fibrosis may not manifest to the same extent in all chambers, the relationship with disease stage remains unknown. As cardiac dysfunction progresses during later stages of disease development, identifying the processes of tissue remodeling across all 4 chambers of the heart in the very early stages of disease development would give critical insight for developing new therapies to prevent cardiac fibrosis in DMD.

Current standard of care for DMD relies primarily on immunosuppression via glucocorticoid administration to prolong ambulatory function and slow disease progression (21). However, glucocorticoids are often accompanied by deleterious side effects, such as attenuated growth, obesity, behavioural disorders, and other detriments (22). The limited cardio-protection offered by emerging gene therapies (23) further underscores the remaining unmet need to develop additional cardioprotective therapies in DMD. In this regard, adiponectin (ApN)-based therapies offer a potential new avenue for consideration given ApN has been shown to be cardioprotective in other models of cardiovascular dysfunction through pleiotropic functions including anti-inflammatory and metabolic-enhancing properties (24,25). Furthermore, the D2.B10-DMD*^mdx^*/2J (D2.*mdx*) mouse model of DMD exhibits decreased plasma ApN levels due to reduced secretion by adipose tissue (26). Transgenic overexpression of ApN in C57BL10.*mdx* mice also improved locomotor activity thus suggesting ApN may have a potential therapeutic role in DMD (26) but the potential to modulate cardiac pathology is unknown. Other beneficial effects of ApN on skeletal muscle in C57BL10.*mdx* mice were observed in models utilizing AdipoRon, which is an orally administered synthetic small-molecule (non-peptide) ApN receptor (AdipoR) agonist (26). Given that ApN turns over quickly and ideally should be injected rather than orally administered to produce its effects (like most large multimeric peptides), ADP355 was pharmacologically developed to overcome these limitations. ADP355, now known as ALY688, is a small peptidomimetic of ApN that activates both AdipoR1 and R2 receptors to induce ApN-like effects, and has exhibited potent anti-fibrotic phenotypes in liver, kidney, and skin fibrosis models (27). It is noteworthy that ALY688 has been developed with pharmacokinetics enabling once-daily injection (unpublished observation).

Recent findings demonstrate that ALY688 preserves certain aspects of mitochondrial bioenergetics in skeletal muscle (28). These effects further support consideration of this compound as a therapeutic intervention for the dystrophin deficient heart. This is important given that mitochondrial stress response pathways represent a potential contributor to cardiac fibrosis development that occurs following a sustained inflammatory response (29). Indeed, reduced mitochondrial pyruvate oxidation and increased mitochondrial reactive oxygen species occurs in the LV of D2.*mdx* at an early age (4-week-old) where fibrosis has yet to manifest (4). Exploring the degree to which adiponectin receptor agonism exerts cardioprotective effects through mitochondrial metabolism in addition to anti-inflammatory and anti-oxidative stress effects is therefore warranted.

The first purpose of this investigation was to examine the degree to which histopathological characteristics develop in all four chambers of the heart in early stages of the disease process in D2.*mdx* mice. The second purpose of the study was to determine the relationship between histopathology, inflammation, and mitochondrial metabolism and whether this differed between the right and left sides of the heart. The final purpose of this investigation was to determine if the adiponectin receptor agonist ALY688 prevented fibrosis and other indices of histopathology in relation to anti-inflammatory effects and mitochondrial reprogramming. We chose to investigate early stages of disease progression by using 4-week-old D2.*mdx* mice given disease symptoms often manifest in young children (30). Our discoveries reveal a robust right ventricular fibrosis and cardiomyocyte hypertrophy as an early event in cardiac remodeling that is defined by mitochondrial-linked apoptosis and metabolic stress as well as a highly specific pro-inflammatory infiltrating macrophage signature that was not seen in the LV. These effects were completely prevented by short-term treatment of the adiponectin receptor agonist ALY688.

## Results

### Fibrosis is dominant in the right ventricle and left atrium in D2.mdx mice and is prevented by ALY688

To determine if fibrosis is present at early stages of disease progression, collagen deposition was assessed by staining frontal (four-chamber) cardiac tissue sections with picrosirius red dye.

Considerable heterogeneity between chambers was noted wherein the RV free wall (epicardial region) and LA of D2.*mdx-*VEH demonstrated increased collagen deposition (7.7-fold and 2-fold increase vs WT, respectively), in contrast to an absence of fibrosis in the LV free wall and RA of D2.*mdx* compared to WT **(Figure 1A, B)**. Both drug doses significantly protected against collagen deposition along the RV free wall (LD, 2-fold decrease vs VEH; HD, 5-fold decrease vs VEH). In the LA, both drug doses significantly protected against collagen deposition (LD, 1.6-fold decrease vs VEH; HD; 3-fold decrease vs VEH) **(Figure 1A, B)**.

**Figure 1.**
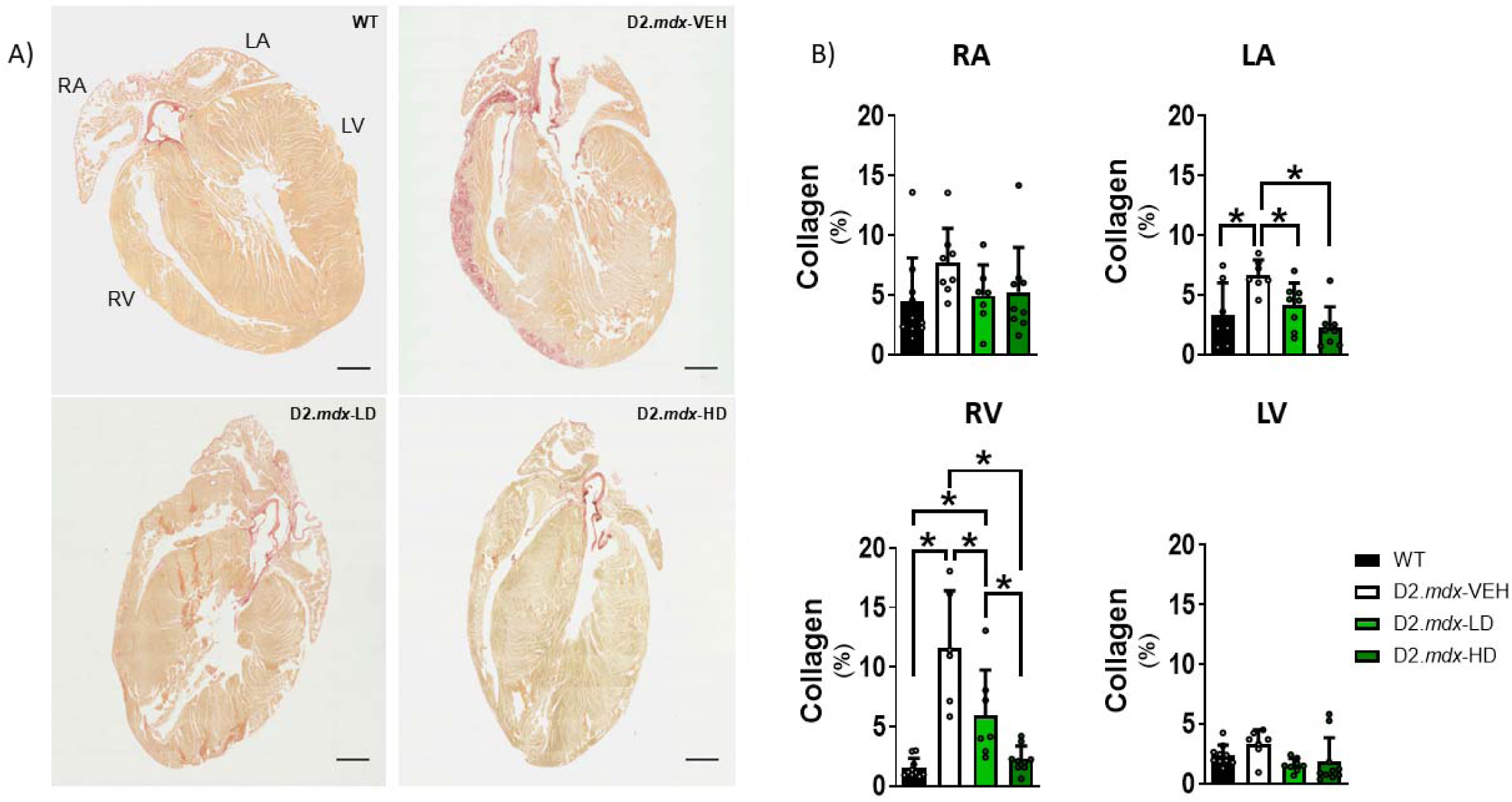
4-week-old D2.*mdx* mice exhibit robust, chamber-specific cardiac fibrosis, which is protected by daily ALY688 administration. A) 5 μM thick frontal sections of paraffin-embedded cardiac tissue were stained with picrosirius red, which selectively binds collagen fibres. B) All four chambers were assessed for collagen deposition. Images were taken with EVOS M7000 Imaging System at 20x magnification. Dark red segments denote collagen. Results represent mean ± SD; n=6-10. Scale bars are 1 mm. All *p* values are FDR-adjusted by Benjamini, Krieger, and Yekutieli *post-hoc* analyses. \**p*<0.05 denotes significance. WT = Wildtype; D2.*mdx*-VEH = vehicle (saline)-treated *mdx*; D2.*mdx*-HD = high dose (ALY688)-treated *mdx*; RA = right atrium; LA = left atrium; RV = right ventricle; LV = left ventricle.

D2.*mdx*-LD mice were excluded from subsequent analyses given that HD was more effective at preventing fibrosis in the affected chambers.

### Right ventricular cardiomyocyte hypertrophy is prevented by ALY688, while myocardial disorganization occurs in a chamber-specific manner in D2.mdx mice

D2.*mdx*-VEH demonstrated larger cardiomyocyte diameter in both ventricles’ vs WT (RV, 16% larger; LV, 15% larger) assessed by average Minimal Feret Diameter indicating that cardiomyocyte hypertrophy may occur early in this model. D2.*mdx*-HD exhibited a complete prevention of this hypertrophy in RV (HD, 14% smaller vs VEH) while MFD remained unchanged in the LV **(Figure 2A)**. When assessing morphometrics between groups from hearts collected over two phases, WT heart weights were significantly larger than both D2.*mdx* groups (VEH, 12% smaller than WT; HD, 14% smaller than WT), however, when normalizing heart weights to tibia length, no significant differences were observed between groups **(Figure 2B)**. D2.*mdx* mice also demonstrated significantly smaller body weights when combining data from both phases (VEH, 25% smaller than WT; HD, 26% smaller than WT) **(Figure 2B)**. Four phases of in-house breeding and daily injections were required to collect chamber-specific data for all measures with a sufficient sample size **(Figure S1A)**. Morphometrics were collected from phase one and four, including ventricle-specific weights during phase four, which were unchanged between groups **(Figure S1B)**.

**Figure 2.**
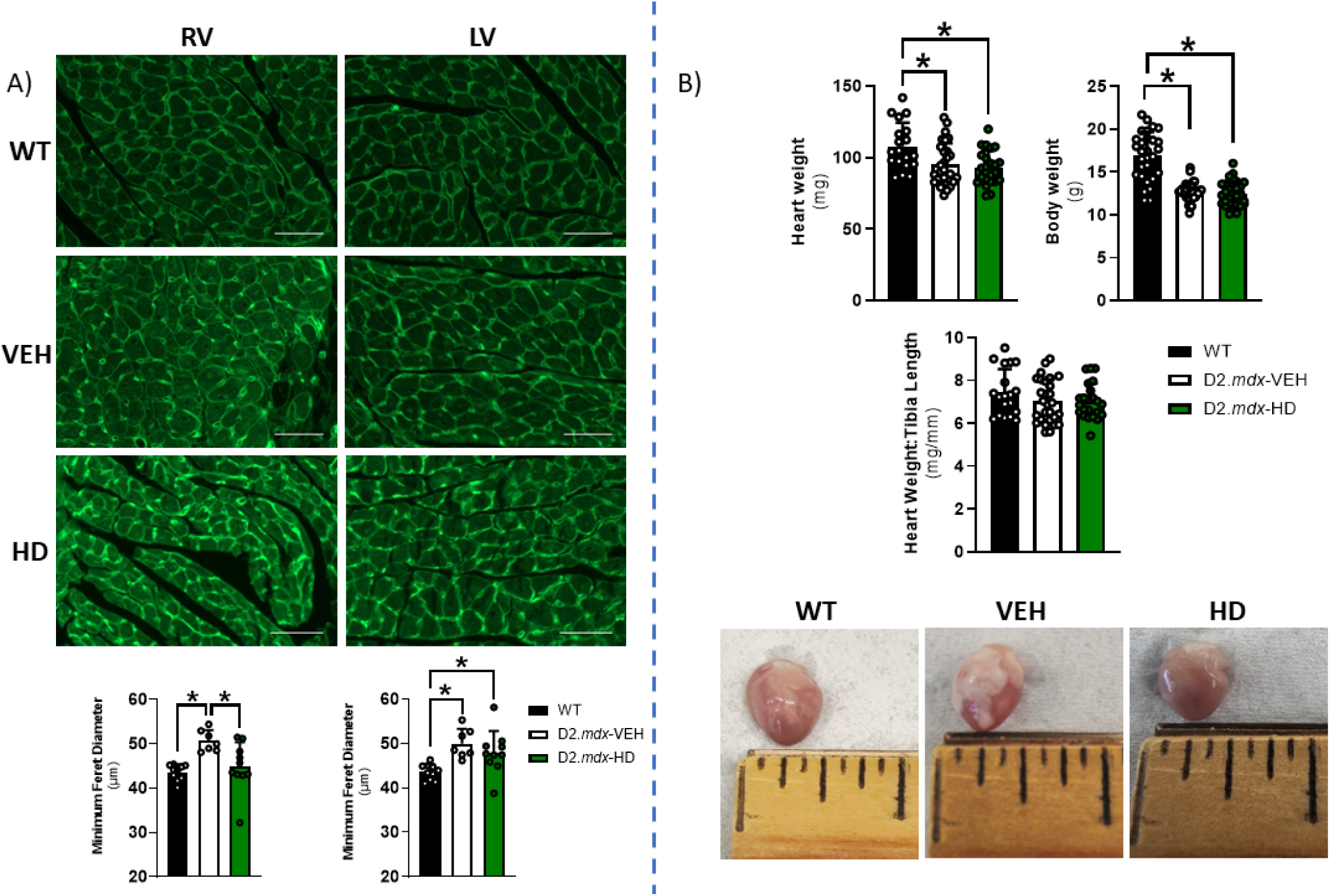
Ventricular cardiomyocytes are hypertrophied in D2.*mdx* mice but rescued by ALY688 in the RV. A) Frontal sections of both ventricles were stained with wheat-germ agglutinin (WGA) and imaged using immunofluorescence. Results represent mean ± SD; n=7-10. Scale bars are 50 μm. All *p* values are FDR-adjusted by Benjamini, Krieger, and Yekutieli *post-hoc* analyses. \**p*<0.05 denotes significance. B) Morphometric data comparing heart weight (mg), body weight (g), and heart weight-to-tibia length (mg/mm) between WT, D2.*mdx*-VEH, and D2.*mdx*-HD, with representative images of perfused whole-heart with the RV oriented frontward. Results represent mean ± SD; n=23-35 (combined phases 1+4). All *p* values are FDR-adjusted by Benjamini, Krieger, and Yekutieli *post-hoc* analyses. \**p*<0.05 denotes significance. WT = Wildtype; VEH = vehicle (saline)-treated *mdx*; HD = high dose (ALY688)-treated *mdx*; RV = right ventricle; LV = left ventricle.

H&E staining was used to identify myocardial disorganization. Loss of nuclear detail and the presence of coalescing nuclei were used as determinants of cardiac muscle damage. In D2.*mdx-*VEH, increased myocardial disorganization was observed in the RV (4.4-fold increase vs WT), LV (1.8-fold increase vs WT), and LA (1.4-fold increase vs WT) **(Figure S2A, B)**. This disorganization was not prevented by ALY688 in any chamber **(Figure S2A, B)**.

### Fibrosis markers are elevated in RV of D2.mdx mice

The RV of D2.*mdx*-VEH demonstrated significant elevations to TGF-β1 (an index of collagen) (30% increase vs WT) **(Figure 3A, B)**. In D2.*mdx*-VEH, α-SMA was significantly elevated in both the RV and RA (RV, 30% increase; RA, 42% increase vs WT). In the RA, D2.*mdx*-HD had elevated α-SMA (36% increase vs WT). In the LA, α-SMA was elevated in both D2.*mdx*-VEH and D2.*mdx*-HD (VEH, 47% increase; HD, 34% increase vs WT), but in the LV, it was only elevated in D2.*mdx*-HD mice (38% increase vs WT) **(Figure S3A, B)**.

**Figure 3.**
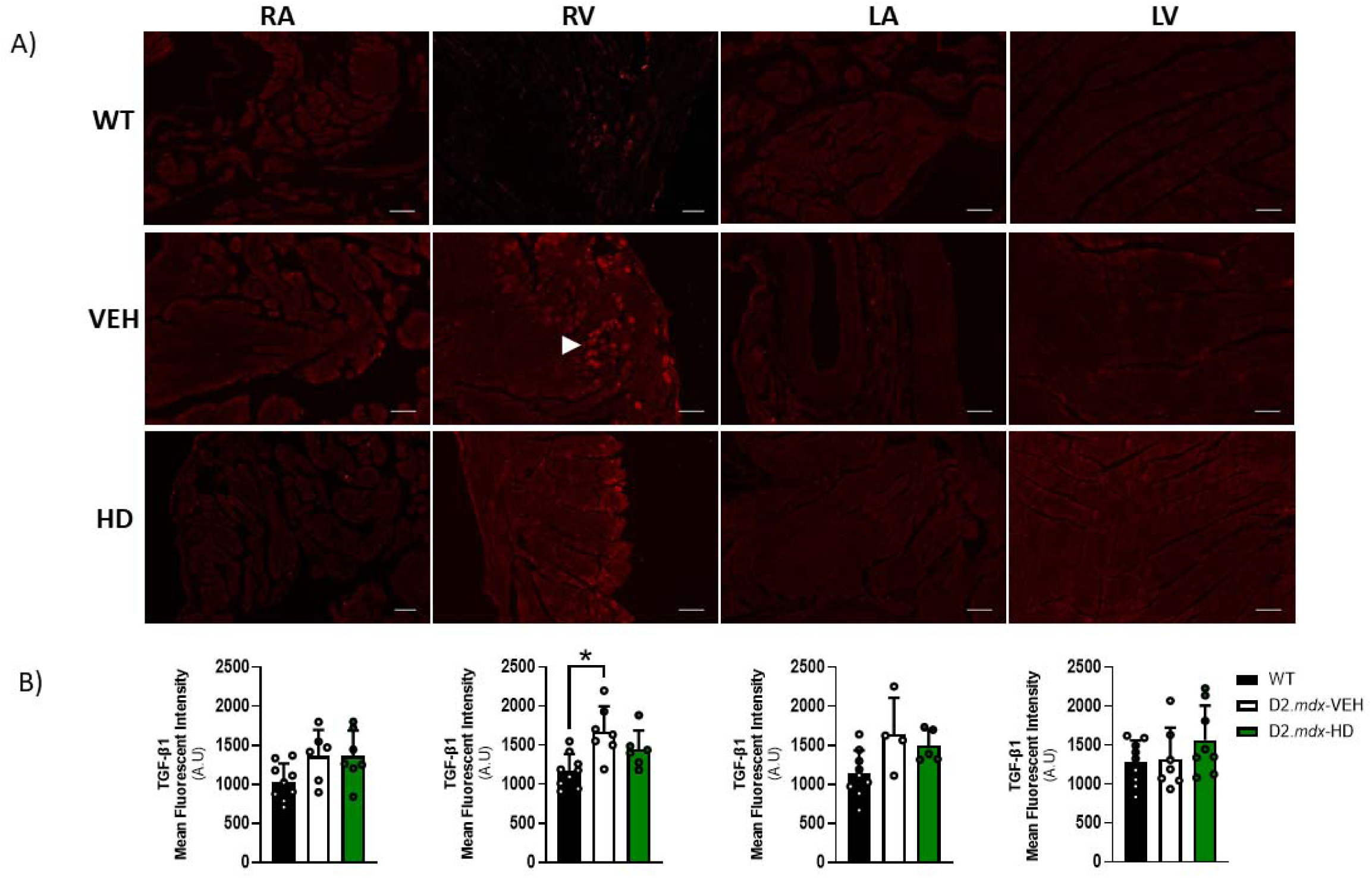
Chamber-specific immunofluorescent staining of TGF-β1. A) TGF-β1 (denoted by white arrows) was quantified by Mean Fluorescent Intensity (A.U.) per region of interest (B). Whole atria and free-wall segments of the ventricles were analyzed. Stains were conducted on 5 μm thick paraffin-embedded frontal sections and imaged by confocal microscopy at 20x magnification. Results represent mean ± SD; n=4-9. Scale bars are 100 μm. All *p* values are FDR-adjusted by Benjamini, Krieger, and Yekutieli *post-hoc* analyses. \**p*<0.05 denotes significance. WT = Wildtype; VEH = vehicle (saline)-treated *mdx*; HD = high dose (ALY688)-treated *mdx*; RA = right atrium; LA = left atrium; RV = right ventricle; LV = left ventricle.

No significant differences between groups were revealed when comparing mean fluorescent intensity of TGF-β1 and α-SMA within the septum **(Figure S4A)**. While only two mice treated with ALY688 demonstrated fibrosis in the RV, we noted reduced markers of both TGF-β1 and α-SMA in the epicardial region **(see representative image, Figure S4B)**.

### Inflammatory cytokines are heterogeneously altered between chambers in D2.mdx

In D2.*mdx-*VEH, the anti-inflammatory cytokine IL-10 was significantly elevated in all four chambers (41-58% increase vs WT). IL-10 was further increased in D2.*mdx*-HD in the LV (31% increase vs VEH) **(Figure S5A, B)**. When assessing the pro-inflammatory cytokine IL-6 across chambers, it was observed that only the LA of D2.*mdx*-HD revealed elevations (31% increase vs WT) **(Figure S6A, B)**.

Both the RA and LA were excluded from remaining measures given not all cryosections could capture the full atria due to their smaller size.

### Spectral flow cytometry reveals significant elevation to an infiltrating macrophage sub-population in RV of D2.mdx-VEH which is protected against in D2.mdx-HD mice

In the RV, CD11b^+^CD45^+^CD64^+^F4/80^+^CCR2^+^ cells, which represent an infiltrating (pro-inflammatory) macrophage population, are elevated in D2.*mdx*-VEH (16-fold increase compared to WT when examining cells normalized per mg tissue), but this is restored back to WT levels in D2.*mdx*-HD **(Figure 4A)**. No differences were detected between groups when examining the same serotype in the LV. When examining CD11b^+^CD45^+^CD64^+^F4/80^+^CCR2^-^ cells, which represent a resident (anti-inflammatory) macrophage population, no differences were detected between groups in either ventricle **(Figure 4B)**. A sample gating scheme retrieved from a D2.*mdx*-HD RV demonstrates how cell populations were carefully selected, or ‘gated’, with special considerations taken for excluding cellular aggregates and debris which may contribute to fluorescent artefact **(Figure S7)**.

**Figure 4.**
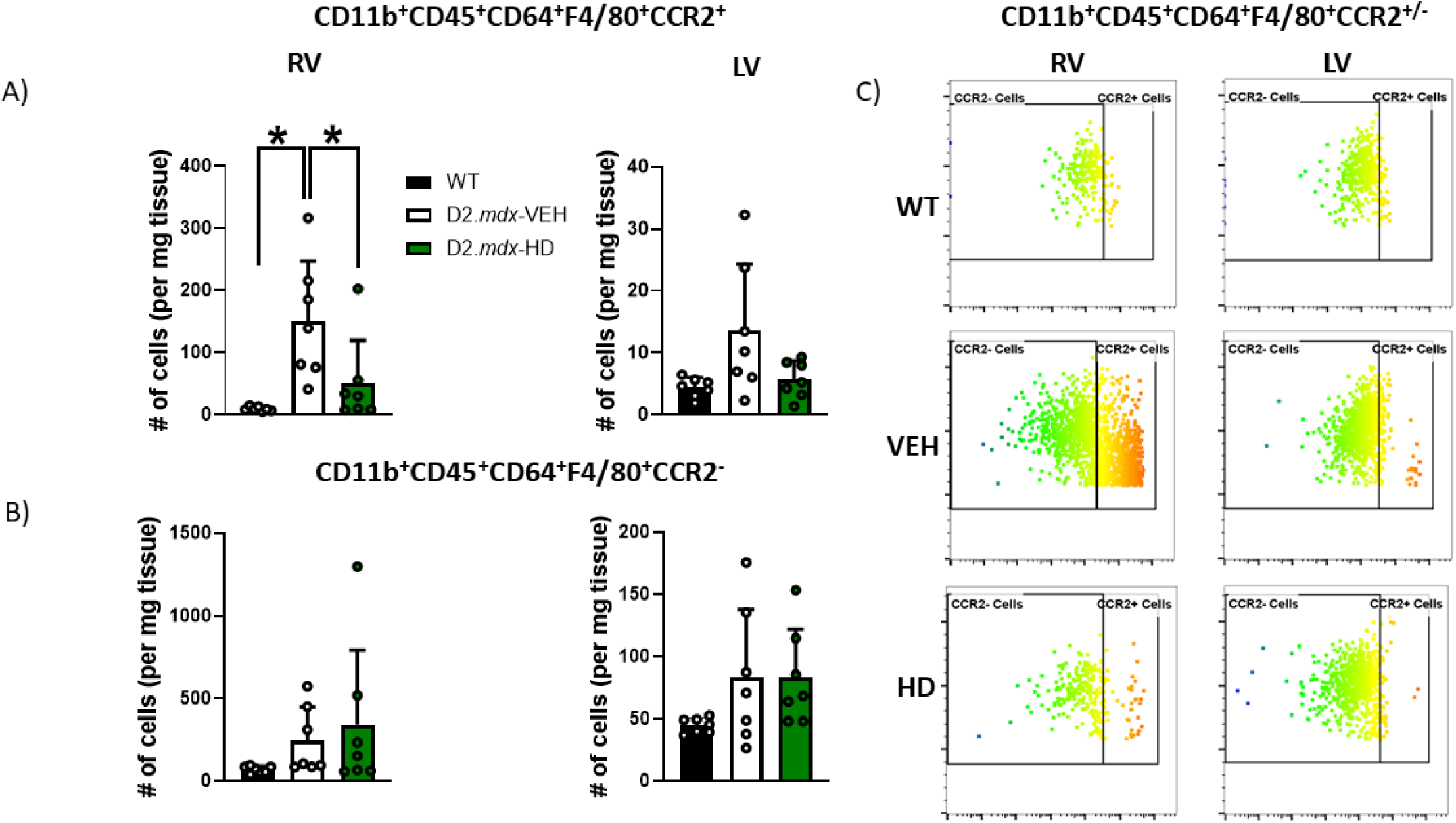
Ventricle-specific spectral flow cytometry analysis of infiltrating and resident macrophage sub-populations in D2*.mdx.* A) Ventricle-specific comparison of CD11b^+^CD45^+^CD64^+^F4/80^+^CCR2^+^ cells (representative of an infiltrating macrophage sub-population), normalized to milligram tissue weight. B) Ventricle-specific comparison of CD11b^+^CD45^+^CD64^+^F4/80^+^CCR2^-^ cells (representative of a resident macrophage sub-population), normalized to milligram tissue weight. C) Representative gating scheme of CCR2^+^/^-^ cells in all three groups between ventricles. Results represent mean ± SD; n=7. All *p* values are FDR-adjusted by Benjamini, Krieger, and Yekutieli *post-hoc* analyses. \**p*<0.05 denotes significance. WT = Wildtype; VEH = vehicle (saline)-treated *mdx*; HD = high dose (ALY688)-treated *mdx*; RV = right ventricle; LV = left ventricle.

### D2.mdx-VEH exhibit indices of mitochondrial stress responses in the RV, but this is only partially restored by ALY688

In the RV, caspase 3 and 9, whose activities are both involved in intrinsic, mitochondrial-mediated apoptotic pathways, are elevated in D2.*mdx*-VEH (caspase 3, 3-fold increase compared to WT; caspase 9, 2-fold increase compared to WT), but are both rescued in D2.*mdx*-HD (caspase 3, 1.7-fold decrease compared to VEH; caspase 9, 1.8-fold decreased compared to VEH) **(Figure 5A)**. No differences were detected in caspase 8 activity, which is involved in extrinsic non-mitochondrial-mediated apoptotic pathways, thereby demonstrating a unique mitochondrial-linked (caspase 9/3) apoptotic signature. In the LV, no differences were detected between groups in any caspase activity.

**Figure 5.**
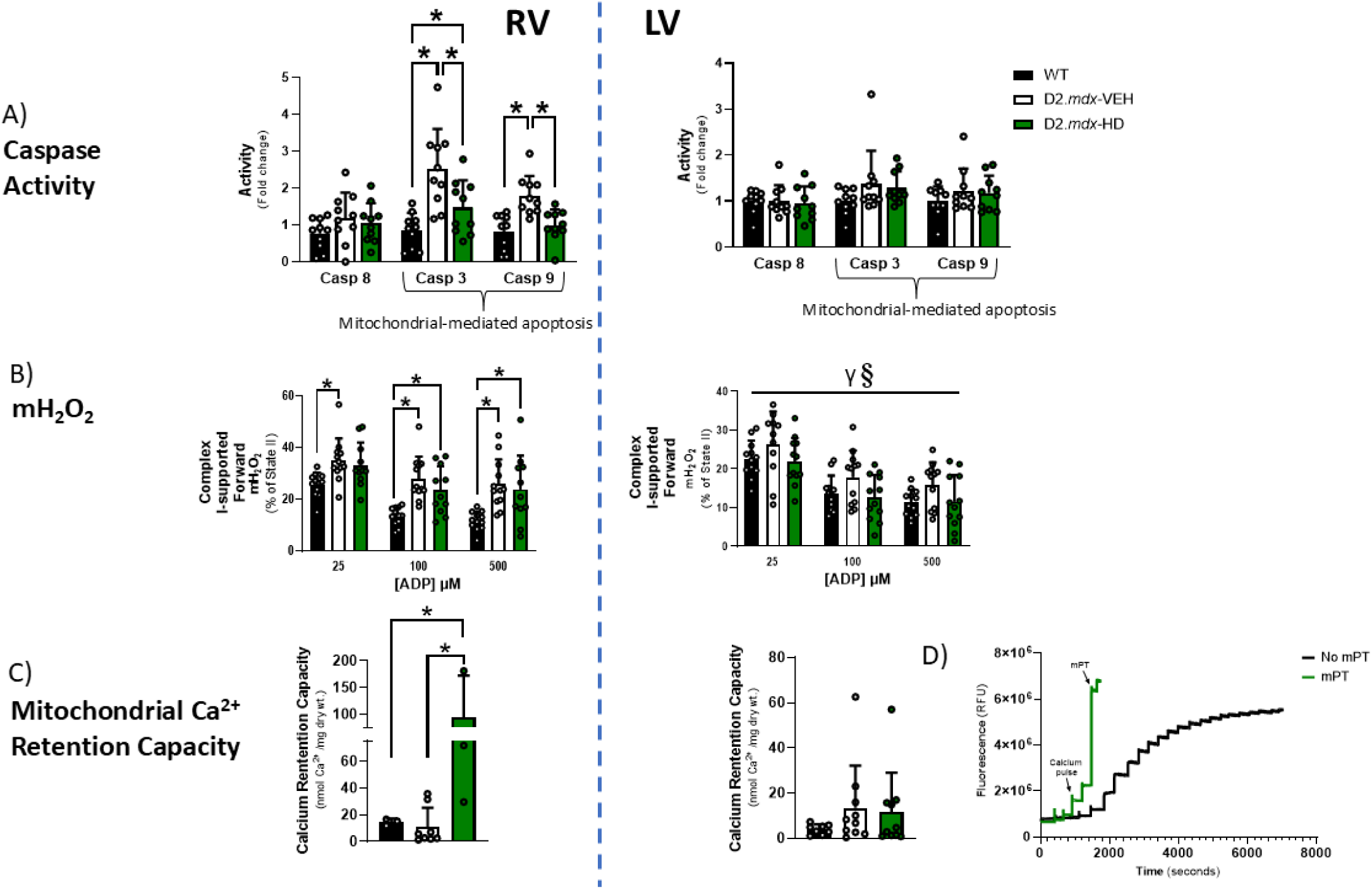
D2.*mdx* exhibit indices of mitochondrial stress primarily in the RV that are partially restored by ALY688 administration. A) Detection of Caspase 3 and 9 activity - representative of intrinsic (mitochondrial-mediated) apoptosis. Caspase 8 activity - involved in extrinsic (non-mitochondrial-mediated) apoptotic pathways - was also examined. Results represent mean ± SD; n=10-12. \**p*<0.05 denotes significance. B) Analysis of mitochondrial complex I-supported (pyruvate + malate) forward electron transfer mH_2_O_2_ (represented as % of State II) in the presence of creatine in both ventricles across metabolic demands (25, 100, 500 μM ADP). Results represent mean ± SD; n=10-12. RV: \**p*<0.05 denotes significance; LV (main effects are denoted by horizontal bar): γ*p*<0.05 WT vs D2*.mdx*-VEH; §*p*<0.05 D2.*mdx*-VEH vs D2.*mdx*-HD. C) Analysis of mitochondrial calcium retention capacity (CRC) to trigger mPT (which precedes mitochondrial-mediated apoptosis). D) Representative CRC trace demonstrating ventricular PmFB with distinct mPT, compared to PmFB without mPT. Results represent mean ± SD; n=3-10. All *p* values are FDR-adjusted by Benjamini, Krieger, and Yekutieli *post-hoc* analyses. \**p*<0.05 denotes significance. Casp = caspase; mH_2_O_2_ = mitochondrial H_2_O_2_ emission; WT = Wildtype; D2.*mdx*-VEH = vehicle (saline)-treated *mdx*; D2.*mdx*-HD = high dose (ALY688)-treated *mdx*; Ca^2+^ = calcium; mPT = mitochondrial permeability transition.

As a complimentary measure to caspase 9/3 activity, mitochondrial calcium retention capacity was examined to determine the ability of mitochondria to uptake calcium prior to triggering permeability transition (mPT), which precedes mitochondrial-mediated apoptosis. In the RV, D2.*mdx*-HD exhibited a significantly elevated capacity to uptake calcium prior to mPT compared to both WT and D2.*mdx*-VEH, which corresponds with caspase 9/3 activity data. However, disease effects were not detectable at this low sample size **(Figure 5C)**. It should be noted that only PmFBs that demonstrated a clear sign of calcium efflux (a sign of mPT) were included. **Figure 5D** demonstrates traces that exhibit mPT versus traces without clear mPT. When pooling traces with a clear opening and traces without clear mPT together, it was determined that D2.*mdx*-VEH had a significantly lower calcium retention capacity compared to WT, which was subsequently rescued in D2.*mdx*-HD **(Figure S8D)**. No differences were detected in the LV using either analytical technique (data reflecting only PmFBs with mPT versus pooled data) indicating that a heightened probability of calcium-induced mPT is unique to the RV.

By titrating pyruvate (5 mM) and malate (2 mM) with 20 mM creatine across increasing submaximal ADP concentrations to model metabolic demands close to physiological conditions (25, 100, and 500 μM) in the RV, it was observed that D2.*mdx-*VEH and D2.*mdx*-HD had elevated mH_2_O_2_ (denoted as a % of State II) (D2*.mdx*-VEH, 27-58% increase vs WT; HD, 28-51% increase vs WT) **(Figure 5B)**. As this measure was performed in the presence of ADP, which stimulates oxidative phosphorylation and attenuates mH_2_O_2_ due to its effects on lowering membrane potential (31) and considering that data were expressed relative to maximal H_2_O_2_, the results indicate that the elevated mH_2_O_2_ in the RV are due specifically to a lower mitochondrial responsiveness to ADP. In the LV, it was observed that D2.*mdx*-VEH had elevated mH_2_O_2_ (14-29% increase vs WT) due to lower mitochondrial responsiveness to ADP’s attenuating effects on mH_2_O_2_, but interestingly, this was protected in D2.*mdx*-HD (17-29% decrease vs D2.*mdx*-VEH) **(Figure 5B)**.

By titrating succinate (10 mM) exclusively (2^nd^ protocol) in the presence of saturating creatine, mH_2_O_2_ (denoted as a % of State II) was decreased in D2.*mdx* compared to WT (Succ, 16% decrease vs D2.*mdx*-VEH; 29% decrease vs D2.*mdx*-HD) with 25 μM ADP. No differences were observed in LV **(Figure S8E)**. Group effects were noted whereby WT mice exhibited significantly elevated mH_2_O_2_ (% of State II) compared to D2.*mdx* across a spectrum of metabolic demands.

### Blunted pyruvate-supported complex I, and complex II, -stimulated respiration in RV of D2.mdx is prevented by ALY688

In the RV, when assessed between groups, mitochondrial oxygen consumption using the NADH-generating substrates pyruvate (5 mM) and malate (2 mM) with saturating (20 mM) creatine was significantly lower in D2.*mdx*-VEH across a spectrum of metabolic demands (25 μM to 5000 μM ADP) (35-53% decrease vs WT) but was protected in D2.*mdx*-HD (36-46% increase vs D2.*mdx-*VEH) **(Figure 6B)**. Similar conclusions were also observed between groups without creatine (D2.*mdx*-VEH, 19-43% decrease vs WT; D2.*mdx*-HD, 7-24% increase vs D2.*mdx-*VEH) **(Figure S8A)** which indicates that mitochondrial creatine-dependent phosphate shuttling may not be modified. The ability of creatine to stimulate respiration was further assessed by comparing respiration rates at submaximal ADP (100, 500 μM) with (20 mM Cr) and without (-Cr) creatine between groups. A complex relationship was identified whereby D2.*mdx*-HD exhibit elevated mitochondrial creatine sensitivity compared to D2.*mdx*-VEH in the RV with 500 μM ADP, while in the LV at 100 μM ADP, creatine sensitivity in D2.*mdx*-HD is significantly decreased compared to both WT and D2.*mdx*-VEH and only significantly decreased compared to WT at 500 μM ADP **(Figure S8B)**. When examining 10 mM glutamate (further NADH generation) and 10 mM succinate (FADH_2_ generation; Complex II stimulation) it was observed that lower mitochondrial oxygen consumption in D2.*mdx*-VEH (glutamate, 52% decrease; succinate, 42% decrease vs WT) was protected in D2.*mdx*-HD (glutamate, 44% increase; succinate, 39% increase vs D2.*mdx*-VEH) **(Figure 6C, D)**.

**Figure 6.**
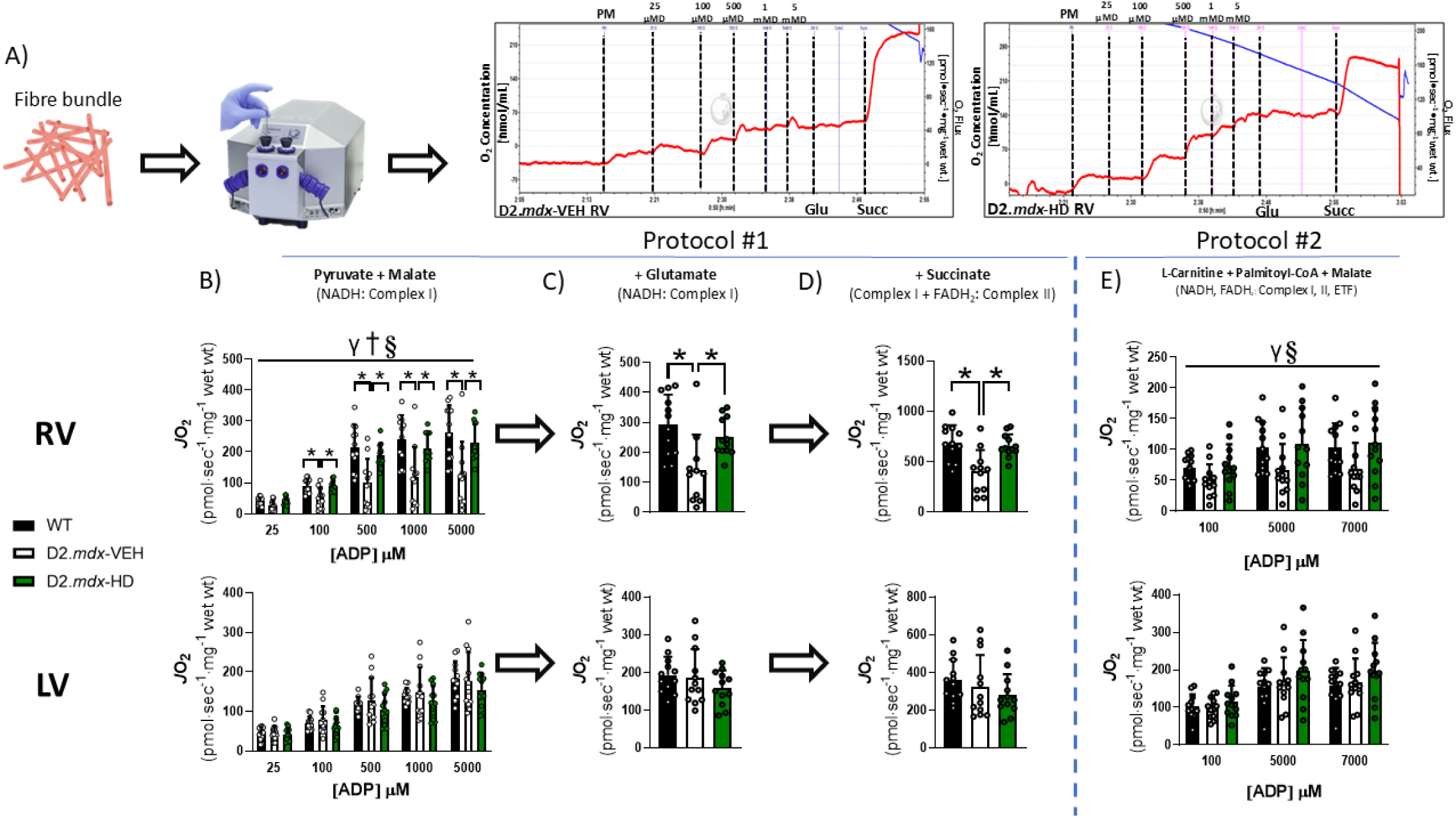
Substrate-specific stimulation of mitochondrial respiration in right and left ventricle of D2.*mdx* mice. A) Representative trace of high-resolution respirometry data demonstrating oxygen consumption that is calculated as the change in slope of oxygen consumption in the respirometry chamber (left trace, VEH RV - pyruvate + malate, ADP titration, glutamate, succinate protocol; right trace, HD RV - pyruvate + malate, ADP titration, glutamate, succinate protocol). B) ADP-stimulated respiration was supported by pyruvate (5 mM) supplemented with malate (2 mM) (NADH, complex-I stimulation) as an index of carbohydrate oxidation. C) After the final addition of ADP (5000 μM) in ‘A’, glutamate (10 mM; NADH, complex-I stimulation) was added as an index of amino acid oxidation, followed by D) succinate (10 mM) (FADH_2_, complex-II stimulation). E) Palmitoyl CoA (20 μM, supplemented with 10 μM L-Carnitine, 500 μM Malate and 100-7000 μM ADP; NADH and FADH_2_) was added to a separate permeabilized fibre bundle as an index of fat oxidation. FADH_2_ from the TCA cycle transfers its electrons at complex-II while FADH_2_ from beta oxidation provides electrons for ETF. NADH = Nicotinamide adenine dinucleotide; FADH_2_ = Flavin adenine dinucleotide; ETF = Electron transport flavoprotein; RV = right ventricle; LV = left ventricle; PM = pyruvate + malate; ‘D’ (from Panel A) = ADP. Results represent mean ± SD, n=10-12; a 2-way ANOVA was used to determine the difference between WT, D2.*mdx*-VEH and D2.*mdx*-HD across ADP concentrations in ‘B’ and ‘E’ while a 1-way ANOVA was used in ‘C’ and ‘D’. All *p* values are FDR-adjusted by Benjamini, Krieger, and Yekutieli *post-hoc* analyses. LV data was log-transformed (for statistical analyses) given that it did not pass parametric testing. \**p*<0.05 denotes significance. Main effects are denoted by horizontal bar over ‘B’ and ‘E’: γ*p*<0.05 WT vs D2*.mdx*-VEH; §*p*<0.05 D2.*mdx*-VEH vs D2.*mdx*-HD; †*p*<0.05 WT vs D2*.mdx*-HD. Oroboros image in Panel A obtained from https://www.oroboros.at/.

No differences were detected in the LV in pyruvate/malate-, glutamate-, or succinate-supported respiration between any groups in the presence of saturating creatine **(Figure 6B)**. In the absence of creatine, differences were only detected in pyruvate/malate-supported respiration in D2.*mdx*-VEH and D2.*mdx*-HD (D2.mdx-VEH, 5-30% increase vs WT; HD, 18-36% increase vs WT) **(Figure S8A)**. In the same protocol, D2.*mdx*-HD demonstrated elevated mitochondrial oxygen consumption (5-21% increase vs D2.*mdx*-VEH) **(Figure S8B)**.

### ALY688 prevents attenuations in fatty acid-supported mitochondrial respiration in D2.mdx

We also examined fatty acid (beta) oxidation in both ventricles by titrating 10 μM L-Carnitine, 20 μM Palmitoyl-CoA, and 500 μM malate, across increasing ADP concentrations, with creatine. In the RV, when assessed between groups, mitochondrial oxygen consumption was lower in D2.*mdx*-VEH (30-39% decrease vs WT) but protected by D2.*mdx*-HD (30-40% increase vs D2.*mdx-*VEH) **(Figure 6E)**.

In the LV, there were no differences in L-carnitine/palmitoyl-CoA/malate-supported respiration between any groups without creatine **(Figure 6E)**.

### Electron transport chain complex-II subunit SDHB protein content may explain improvements to respiration in D2.mdx-HD

Collectively, the lower respiration under all substrate conditions seen in the RV of the disease group, and the preservation effect of the drug, suggests that a common feature of these metabolic pathways may be universally modified. In order to gain more insight into this pattern, we recognized that the normalization of respiration per mg of tissue could be influenced by potential shifts in the contents of electron transport chain proteins, particularly those that are common and downstream of Complex I and II stimulation.

However, no changes in subunits of Complex III, IV or V (ATP Synthase) were seen in the RV. Of interest, complex-II subunit SDHB was significantly elevated in D2.*mdx-*HD (16% increase vs D2.*mdx-*VEH) which may have contributed to the higher succinate-stimulated respiration in this group. No changes in the measured ETC proteins were observed in the LV. Total ETC content was also pooled together, but there were no differences between groups in either ventricle **(Figure S8C)**.

### Mitochondrial signaling and mitophagy protein content does not explain divergent bioenergetic responses in the RV

We examined mtCK due to its involvement with ATP/ADP cycling, however, no significant differences were observed between groups in either ventricle **(Figure 7A)** which is consistent with the observations that changes in respiration in disease and drug-treated groups were similar whether creatine was present or absent in the assay. Collectively, this suggests that creatine-dependent phosphate shuttling was not modified by either group. Total and phosphorylated protein content of AMPK were also examined, which are thought to be activated by adiponectin-receptor agonism (32). In the RV, both D2.*mdx-*VEH and D2.*mdx-*HD mice had elevated phosphorylated AMPK (D2.*mdx-*VEH, 21% increase vs WT; HD, 27% increase vs WT; **(Figure 7A)**. In the LV, D2.*mdx-*VEH mice demonstrated elevated phosphorylated AMPK (32% increase vs WT) which was prevented by the drug **(Figure 7A)**. Lastly, the redox-sensitive apoptosis-activating protein p38 MAPK, which can be phosphorylated in response to mH_2_O_2_ (33) and activated by adiponectin as demonstrated in other studies (34–36), was examined to determine if it was altered between groups (33,37). No differences were detected in total or phosphorylated p38 MAPK between groups in either ventricle **(Figure 7A)**.

**Figure 7.**
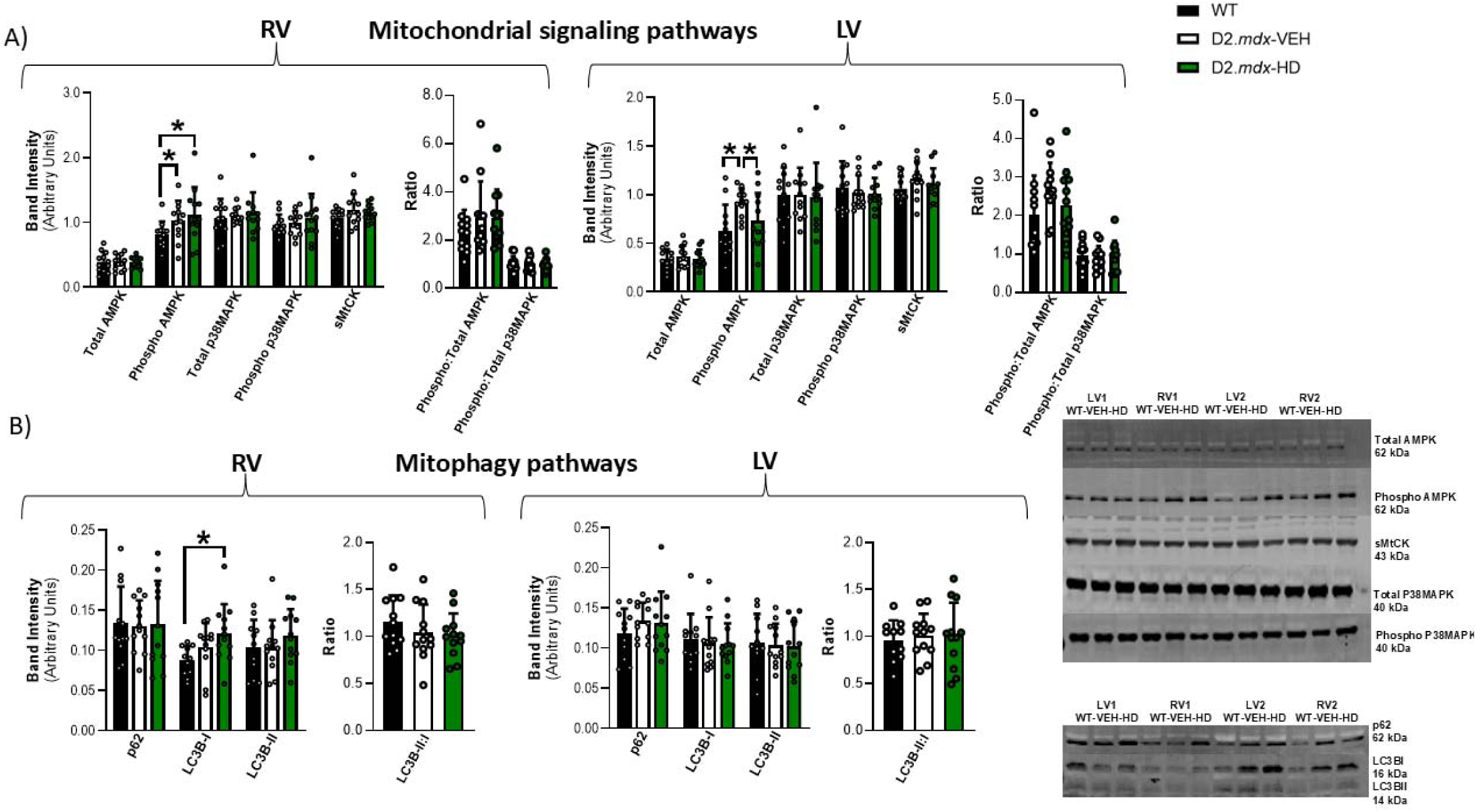
Western blots of mitochondrial signaling and mitophagy markers. (A) Ventricle-specific mitochondrial signaling markers. (B) Ventricle-specific mitophagic markers. Results represent mean ± SD; n=11-12. All *p* values are FDR-adjusted by Benjamini, Krieger, and Yekutieli *post-hoc* analyses. The ‘ratio’ is defined as phosphorylated protein content divided by total protein content. \**p*<0.05 denotes significance. RV = right ventricle; LV = left ventricle.

**Figure 8.**
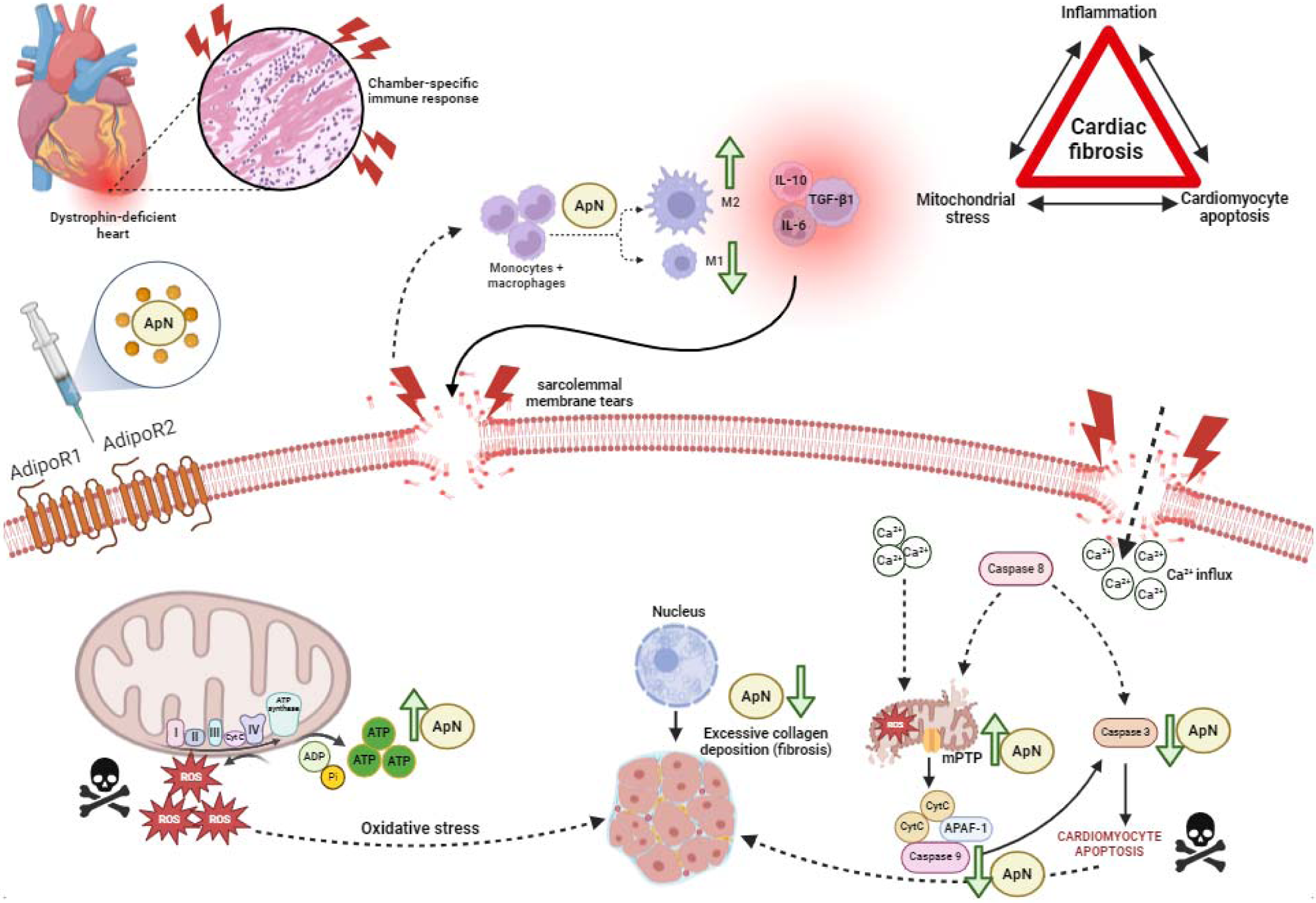
Proposed mechanisms underlying dystrophin deficiency-induced cardiac fibrosis in 4-week-old D2.*mdx* mice that can be partially rescued by daily adiponectin-receptor agonism. We establish that three major secondary contributors to dystrophin deficiency-induced cardiac fibrosis (inflammation, mitochondrial-mediated apoptosis, and mitochondrial stress) differentially impact chambers of the heart. Our findings indicate that these secondary contributors demonstrate rescue effects when D2.*mdx* mice are injected daily with ALY688, ultimately leading to reduced cardiac fibrosis in the right ventricle (RV). To examine indices of mitochondrial stress, we investigated chamber-specific mitochondrial respiration and mH_2_O_2_, while intrinsic (mitochondrial-mediated) apoptosis was assessed by a combination of calcium retention capacity and caspase activity as complimentary measures. Inflammation was assessed by spectral flow cytometry to determine shifts in macrophage polarization (infiltrating versus resident macrophage sub-populations) as well as by immunofluorescence of various inflammatory and fibrosis-linked cytokines. Our results indicate that daily adiponectin-receptor agonism may represent a viable therapeutic intervention for rescuing RV fibrosis in young D2.*mdx* mice. Green arrows denote protective effects of ALY688 observed in the D2.*mdx*-HD RV across various remodeling signatures, including shifts to pro-/anti-inflammatory markers, mitochondrial stress responses, and cardiac fibrosis.

Mitophagy serves to preserve mitochondrial function in cardiac tissue. Impairments to mitophagic processes can induce mitochondrial dysfunction, which can exacerbate cardiomyopathy (38). Protein content of the autophagosome-ubiquitination adaptor proteins SQSTM1/p62 and LC3BII/I were investigated, however, the only difference detected was in LC3B-I, where D2.*mdx*-HD was significantly elevated compared to WT in the RV **(Figure 7B)**.

## Discussion

Cardiac fibrosis is a hallmark feature of cardiomyopathy in DMD yet there is no therapy proven to delay its progression. Furthermore, the degree to which fibrosis and related cellular pathologies occur across each chamber of the heart remains unclear, particularly in the early stages of disease, due in part to the limited tissue mass available for comprehensive tissue assessments in pre-clinical models. To overcome this challenge, we implemented a considerable breeding strategy to quadruple the number of hearts available for comparative tissue assessments (**Figure S1A**). With this approach, we employed a comprehensive 4-chamber assessment of histopathology in the hearts of very young dystrophin-deficient mice to capture the early stages of disease progression. The discovery that fibrosis is dominant in the RV and LA demonstrates that dystrophin deficiency causes chamber-specific histopathology in the heart. The dominant RV fibrosis, and an accompanying cardiomyocyte hypertrophy, were linked to a pro-inflammatory signature defined by a highly specific macrophage sub-population that has not been reported previously in DMD. Mitochondrial-linked apoptosis and reduced pyruvate oxidation further defined a RV pathology not seen in the LV. We also show that a peptidomimetic adiponectin receptor agonist completely prevents RV fibrosis, cardiomyocyte hypertrophy, inflammation, and mitochondrial stress as well as LA fibrosis which opens a new avenue for exploring the potential for anti-fibrotics to treat cardiomyopathy in DMD.

### Right ventricular and left atrial fibrosis are dominant features of very early-stage cardiac histopathology in young D2.mdx mice

Our discovery of a robust RV epicardial and LA fibrosis at only 4 weeks of age in D2.*mdx* mice is notable given no fibrosis was detected in the RA or LV. RV and LA fibrosis are therefore unique hallmarks of the very early stages of cardiomyopathy in D2.*mdx* mice. Cellular disorganization, indicative of damage and mononuclear cell infiltrate, was greatest in the RV but also present in LA and LV, while both ventricles demonstrated cardiomyocyte hypertrophy. Through this pattern, a consistent dominant pathology in the RV could be seen. Flow cytometry revealed a greater content of infiltrating CD11b^+^CD45^+^CD64^+^F4/80^+^CCR2^+^ macrophages (referred to as CCR2^+^) in the RV that was not elevated in the LV consistent with the lack of fibrosis detected in this chamber. As resident CD11b^+^CD45^+^CD64^+^F4/80^+^CCR2^-^ (referred to as CCR2^-^) macrophages were not altered in the RV, these findings position infiltrating macrophages as a distinct inflammatory signature of early-stage RV fibrosis in DMD. This finding opens a new avenue for research to understand how this specific macrophage sup-population mediates early chamber-specific remodeling and eventual cardiac dysfunction in DMD.

The cause of heterogeneous tissue pathology across the heart is unclear but perhaps intuitive given each chamber is defined by their unique features such as wall thickness, extracellular matrix regulation (39,40) as well as metabolic and redox phenotypes (41,42), pressure/load differences, amongst other attributes. It is particularly notable that fibrosis occurred only in chambers that are directly connected to the pulmonary circulation (RV, LA). While the cause for this pattern is unclear, the findings raise questions regarding pulmonary arterial pressures, particularly regarding RV fibrosis. For example, RV fibrosis and hypertrophy occurs during pulmonary hypertension in other conditions (43–45) and pulmonary hypertension was also identified in a small sample of people with DMD (46,47). Separate studies reported RV fibrosis in some but not all post-mortem case reports (12,19) and in later stages of disease in adult C57BL/10ScSn-Dmd*^mdx^* mice (48,49). RV hypertrophy was also reported in people with DMD (50) and 11-month-old C57BL/10-*Dmd^mdx^*subjected to exercise (49). While no study has performed concurrent measures of RV fibrosis, hypertrophy and pulmonary arterial pressures, RV epicardial fibrosis and increased pulmonary vascular resistance were found in adult 16–17-week-old D2.*mdx* mice (51) suggestive of possible pulmonary arterial hypertension. Likewise, a recent case report in an adult with DMD measured increased filling pressures in both RV and LV (47) with elevated pulmonary artery pressure that was related to respiratory difficulty and a reduced ability to cough. RV arrhythmias and systolic dysfunction have also been reported in children and adults with DMD (52,53) that could explain elevations in RA pressure (46). These separate observations of respiratory insufficiencies, pulmonary hypertension and RV pathology align with a previous proposal that respiratory muscle weakness may drive RV pathology in DMD (48). We have also previously reported considerable diaphragm weakness and reduced inspiratory pressures during tracheal occlusion (28,54,55) at the same young age of D2.*mdx* mice used in the current study that found RV fibrosis. In keeping with a link to pulmonary circulation, the LA fibrosis we observed could be examined in relation to pulmonary venous pressures and atrial ECG given atrial fibrillation has been linked to LA fibrosis in certain conditions (56). Atrial fibrillation has also been observed in people with DMD, although this occurred in patients demonstrating a LV ejection fraction of <35% (57), as has LA fibrosis (20). As we previously reported no change in ejection fraction in 4-week-old D2.*mdx* mice (4), it is likely that the RV and LA fibrosis seen in the present study precede an eventual dysfunction that could be re-examined through a time-course comparison throughout disease progression and in relation to pulmonary circulation and the well-established respiratory muscle weakness in the D2.*mdx* model.

### The adiponectin receptor agonist ALY688 prevents chamber-specific fibrosis and infiltrating macrophage accumulation

The RV and LA fibrosis in D2.*mdx* mice were partially and completely prevented by low and high doses of ALY688, respectively. While we did not have a standard-of-care positive control such as glucocorticoids, to our knowledge no study has ever reported a complete prevention of cardiac fibrosis with this front-line medication. ALY688 also completely prevented RV cardiac fibre hypertrophy. Of interest, ALY688 did not prevent LV cardiomyocyte hypertrophy nor disorganization/damage detected by H&E stains in RV, LA or LV. This finding is consistent with our previous report showing lower diaphragm fibrosis without changes in H&E in response to ALY688 (28). The current findings highlight an interesting disconnect between fibrosis and damage, at least in the therapy-responsive RV and LA, which may otherwise be expected to occur in tandem. Nevertheless, the prevention of fibrosis and cardiomyocyte hypertrophy in the RV is profound given the clinical and pre-clinical relationships between fibrosis and RV dysfunction seen in past literature as discussed in the previous section. The findings serve as a foundation to determine whether the eventual development of RV dysfunction in later stages in D2.*mdx* mice are prevented by this early anti-fibrotic and anti-hypertrophic effect.

Both adiponectin and ALY688 possess anti-inflammatory properties (58–60) but the precise mechanisms remain unresolved. Here, we discovered high dose ALY688 prevents the increase in RV CD11b^+^CD45^+^CD64^+^F4/80^+^CCR2^+^ infiltrating macrophages seen in D2.*mdx* vehicle treated mice. CCR2 signaling is of particular interest in dystrophin deficient mouse models given that previous literature has demonstrated elevations to CCR2 expression and its chemokine ligands, alongside macrophages excessively skewing towards a proinflammatory phenotype in *mdx* diaphragms (61). Additionally, CD11b^+^CD45^+^CD64^+^F4/80^+^CCR2^-^ resident macrophage contents in either ventricle were not affected by ALY688, nor by the disease state as mentioned above. Collectively, we propose that RV disorganization and damage during DMD leads to a unique infiltrating macrophage inflammatory signature that mediates RV epicardial fibrosis given this relationship was completely prevented by ALY688. This CCR2^+^ infiltrating macrophage signature is more directly related to fibrosis than damage given the LV showed increased H&E staining without fibrosis while CCR2^+^ infiltrating macrophages were unchanged. This comparison between both ventricles underscores the distinct fibrotic relationship with CCR2^+^ infiltrating macrophages and their high responsiveness to adiponectin receptor agonism which has not previously been reported.

Immunofluorescent detection of the cytokines IL-6, IL-10, and TGF-β as well as the mature myofibroblast marker α-SMA (62–65), a downstream target of TGF-β, did not reveal consistent patterns in relation to chamber-specific fibrosis. While RV and LA α-SMA were increased in D2.*mdx* vehicle treated mice similar to fibrosis, its content was also elevated in the RA despite no sign of fibrosis in this chamber. IL-10 is thought to be an anti-inflammatory cytokine in DMD (66) yet it was also elevated in the fibrotic RV and LA which might be indicative of a counterresponse to inflammation. TGF-β is a master regulator of fibrosis (67) and was elevated in dystrophic RV consistent with fibrosis in this chamber, yet it was not altered in any other chamber. Also, none of these measures showed a definitive pattern with ALY688’s anti-fibrotic effects in RV or LA. Collectively, immunofluorescent detection of cytokines is insufficient to understand the mechanisms regulating inflammation and fibrosis in 4-week-old D2.*mdx* mice which underscores the importance of the flow cytometry-based detection of RV CCR2^+^ infiltrating macrophages noted above.

### ALY688 prevents RV-specific mitochondrial pro-apoptotic and attenuations in substrate oxidation

Dystrophin deficient muscle, including the heart, causes increased influx of calcium from the extracellular environment partially through damage to the cell membrane (68,69). Calcium overload triggers mitochondrial swelling and reactive oxygen species production leading to formation of a pore across the double mitochondrial membrane system (70). A subsequent event known as ‘mitochondrial permeability transition’ (mPT) activates apoptosis whereby efflux of excess mitochondrial calcium and other pro-apoptotic factors into the cytoplasm triggers caspase 9 which then activates the effector caspase 3 that cleaves numerous essential functional and structural proteins (71). We discovered that RV fibrosis in D2.*mdx* mice was associated with elevated caspase 9 and 3 activities while mitochondrial-independent caspase 8 was not affected. Remarkably, ALY688 completely prevented the increases in caspase 9 and 3 activities. This drug effect was linked to a lower mitochondrial sensitivity to calcium-induced mPT upstream of these caspases indicated by a greater calcium retention capacity prior to excess calcium release. Unlike the RV, there was no disease or drug effect on LV mitochondrial-linked apoptosis consistent with the lack of fibrosis in these groups compared to wildtype. As the degree of damage in LV from D2.*mdx*-vehicle treated mice was about 1/3^rd^ of the level in RV, the data suggests a very high level of damage is required to trigger mitochondrial-linked apoptosis. Overall, the findings demonstrate that adiponectin receptor agonism regulates mitochondrial-linked apoptosis which is prevented by ALY688.

RV fibrosis was also related to attenuated pyruvate (carbohydrate) and fatty acid-stimulated respiration that was prevented by ALY688. This mitochondrial signature is unique to the RV as no changes were seen in the LV. These RV mitochondrial responses were due to altered intrinsic properties of mitochondria given there were no changes in the contents of mitochondrial electron transport chain proteins that were stimulated during the respiration protocols. The mechanistic relationship between attenuated substrate oxidation and RV fibrosis is conceptually difficult to propose as it seems unlikely that lower mitochondrial ATP synthesis would directly cause cell death or modulate the process of fibrosis given alternative energy sources such as from glycolysis would need to be considered in such theoretical models. Nevertheless, this finding may be a sign of a grander mitochondrial stress consistent with calcium overload in DMD as implicated by the mitochondrial-linked apoptosis results. However, ALY688 did not prevent increased in RV mH_2_O_2_ assessed *in vitro* despite doing so in the LV which did not demonstrate fibrosis. While adiponectin receptor signaling is not known to modulate mitochondrial reactive oxygen species, the pattern does not demonstrate a relationship between mH_2_O_2_ and fibrosis in the dystrophic heart although targeted mitochondrial antioxidant compounds would be required to explore this relationship more fully.

Mitophagic markers like p62 and LC3BII/I were both investigated to determine if the cell’s inherent ability to eliminate dysfunctional mitochondria was disrupted. Previous reports in C57BL/10 *mdx* mice demonstrate either increased (68,69) or decreased markers of mitophagy (72,73). Here, we found untreated D2.*mdx* mice did not demonstrate changes in these markers, although ALY688 increased LC3B-I which is upstream of LC3B-II. As no changes in the ratio of these proteins were noted, it is unclear if ALY688 altered mitophagy in a way that could explain the preservation of RV mitochondrial bioenergetics.

Collectively, the data supports a model whereby RV disorganization and damage, as shown by H&E stains, arising from dystrophin mutations causes calcium-induced mitochondrial permeability transition and apoptosis as reviewed previously (69) that leads to a distinct inflammatory state defined by CCR2^+^ infiltrating macrophages in very early stages of disease.

### Perspectives, limitations and conclusions

While cardiac function was not assessed in this study, we have previously reported no change in left ventricular ejection fraction or cardiac output in this very young age of 4-weeks in D2.*mdx* mice using echocardiography in anaesthetized mice (4). This finding does not rule out dysfunctions at 4-weeks that may be detected with stress tests such as dobutamine which was used in older C57BL/10ScSn-Dmd*^mdx^* mice to reveal reduced RV and LV diastolic and systolic functions (48). However, the lack of overt cardiac dysfunction measured in left ventricles by echocardiography in a non-stimulated state seen in our prior work (4) is reflective of what mice experience day-to-day in the absence of any measurable form of stress and is directly related to the histopathological observations reported herein. Therefore, the RV and LA fibrosis as well as ventricular cardiomyocyte hypertrophy seen in the present study very likely precedes an eventual cardiac dysfunction at later stages of the disease as reported previously in D2.*mdx* mice (2). The very small RV of these mice also prevented the use of echocardiography and invasive hemodynamics unlike what is possible in older mice for this chamber (48). These challenges are inherent in assessing very early-stage disease processes in *mdx* mice which was the focus of this investigation. Nonetheless, our findings serve as a foundation to explore the potential for early intervention with ALY688 to prevent cardiac fibrosis and whether this delays the onset of dysfunction in specific chambers including arrythmias given fibrosis is a well-established cause of this anomaly.

The effects of ALY688 were not related to changes in AMPK or p38MAPK which are known to be regulated by adiponectin receptors (74,75). It is possible that the 24-hour gap between the last injection and tissue removal missed the activation of these cascades given that the half life of ALY688 in rodent models is less than this time frame (unpublished data from Allysta). Also, despite using an extensive 4 phase design of breeding to increase tissue availability of all 4 chambers, we were still limited by tissue mass to perform additional measures such as mitochondrial morphology, gene expression-based measures of cytokine dynamics, tissue calcification, and other measures. In this regard, the present findings provide extensive insight considering the limited tissue availability of specific chambers.

Many emerging gene therapies such as exon skipping strategies hold promise to treat DMD. This approach requires considerable research given the need to customize therapies to each of the 70 known dystrophin mutations. Developing additional therapeutic paradigms that target secondary contributors to cardiomyopathy could benefit most people with DMD regardless of the underlying mutation. In this way, adiponectin receptor agonism opens a new paradigm for continued therapeutic development in DMD.

In summary, we discovered a distinct RV and LA fibrotic relationship in early-stage disease of D2.*mdx* dystrophin-deficient mice that occurred in the absence of LV fibrosis. A unique CCR2^+^ infiltrating macrophage sub-population and mitochondrial pro-apoptotic signature defined early-stage RV fibrosis and cardiomyocyte hypertrophy. Remarkably, the adiponectin receptor agonist ALY688 completely prevents RV fibrosis, cardiomyocyte hypertrophy, mitochondrial stress, and CCR2^+^ infiltrating macrophages. As these events occurred at a disease stage preceding overt cardiac dysfunction (4), the discoveries open a new avenue of research to understand the precise mechanisms by which CCR2^+^ macrophages and mitochondrial remodeling contribute to fibrosis, and whether prevention of this relationship with RV histopathology can prevent the eventual decline in heart function. Moreover, the intriguing relationships between RV and LA fibrosis reveal an opportunity to examine the potential role of altered pulmonary circulation in the context of how dystrophin deficiency impacts the heart indirectly by causing weakness in non-cardiac muscles such as the diaphragm. Pursuing this potential integrated paradigm of cardiorespiratory dysfunction with unique chamber-specific inflammatory and metabolic signatures could hold new promise into developing therapies that prevent cardiac fibrosis and dysfunction in DMD.

## Materials and methods

### Animal care

Male D2.*mdx* mice were bred from an in-house colony established at York University’s rodent facility (Toronto, Ontario). Mice were housed with littermates in standard vivarium cages and aged to four weeks until sacrifice (aged 28-32 days), in a room maintained on a 12:12-h light-dark cycle with *ad libitum* access to standard chow and water. Due to low in-house breeding success, 3-week-old DBA/2J Wildtype (WT) mice were ordered directly from Jackson Laboratories (Bar Harbor, USA) and allowed to acclimate for one week prior to sacrifice. Experiments, procedures, and personnel were all approved in accordance with both the Animal Care Committee at York University (AUP Approval Number 2016-18) and with the Canadian Council on Animal Care.

### Experimental design, breeding regimen, and ALY688 drug treatment

Two groups of D2.*mdx* mice were treated with ALY688 (Allysta Pharmaceuticals) formulated at different dosages: low dose (LD; 3 mg/kg body weight/day) and high dose (HD; 15 mg/kg body weight/day) – both of which were dissolved in saline. A separate group of D2.*mdx* mice (VEH, vehicle) were treated with saline. Mice were treated starting at 7-days of age (until 28-32 days of age) via subscapular subcutaneous injection at a dosage of 1 μL/g body weight. WT mice did not receive injections.

Due to limited tissues in each chamber of the heart, four mice per ‘n’ were needed to generate all data (**Figure S1A)**. Accordingly, four breeding phases of D2.*mdx* mice were required to obtain sufficient tissue for all analyses. During phase 1, hearts from each group of mice were processed for tissue histology and immunofluorescence. Given that LD and HD yielded similar fibrotic responses, as assessed by picrosirius red staining, it was decided that LD would not be pursued for analyses beyond phase 1. During phase 2, hearts from each group of mice (now WT, D2.*mdx-*VEH, and D2.*mdx-*HD) were flash frozen for Western blot analyses. During phase 3, hearts from each group of mice were incubated in preservation buffer and subsequently processed for mitochondrial bioenergetic assays. Phase 4 was conducted to obtain additional morphometric data, additional mitochondrial bioenergetics data, caspase activities, and spectral flow cytometry data. Sample sizes are indicated in figure legends where appropriate.

### Surgical procedure

Hearts were carefully excised while the animal was anesthetized with medical-grade isoflurane. For phase 1, whole-hearts were preserved in 10% neutral-buffered formalin (Millipore Sigma, Burlington, MA, USA) and processed for histology. Morphometric data such as heart weight was also collected during this phase. For phase 2, hearts were carefully separated into 4 chambers under a microscope with surgical-grade scissors and were then frozen in liquid N_2_ and stored at -80°C for Western blots. For phase 3, ventricles were removed and placed in ice-cold BIOPS preservation buffer containing (in mM) 50 MES Hydrate, 7.23 K_2_EGTA, 2.77 CaK_2_EGTA, 20 imidazole, 0.5 dithiothreitol, 20 taurine, 5.77 ATP, 15 PCr, and 6.56 MgCl_2_•6H_2_O (pH 7.1), before being separated into fibre bundles for mitochondrial bioenergetic assays (see below). During phase 4, hearts with either placed in ice-cold BIOPS preservation buffer or processed for spectral flow cytometry.

### Histological and immunofluorescent staining

#### Preparation of microscope slides

Hearts were immediately submerged into a 15 mL canonical Falcon tube containing 10% neutral-buffered formalin and stored for 24 hours at room temperature. Following a 24-hour incubation period in formalin, hearts were transferred to a separate 15 mL conical centrifuge tube containing 70% ethanol and stored at 4°C thereafter.

Once ready for tissue processing, hearts were individually placed in plastic paraffin-embedding cassettes (Simport Scientific, Saint-Mathieu-de-Beloeil, QC) where they were dehydrated using a gradient of ethanol concentrations (70% to 99%), followed by two 100% xylene incubations. Cassettes were placed overnight in an oven heated to 54-57°C, which contained melting Type H paraffin wax (Thermo Fisher Scientific, USA), in a process intended to allow paraffin wax to penetrate and embed the samples. The following morning, after cassettes were removed from the oven and the paraffin wax had solidified at room temperature, paraffin within the cassettes was again heated to melting point, and samples were transferred to disposable base molds. Following this, paraffin was once again re-introduced to the cassettes and left to solidify at room temperature before the samples were ready to be sectioned at 5 μM thickness on a frontal (4-chamber) plane using a microtome and hot water bath. Finally, sections were carefully placed onto Fisherbrand SuperFrost Plus positively charged microscope slides (Thermo Fisher Scientific) for microscopy.

#### Picrosirius Red (PSR)

Fibrosis was assessed on formalin-fixed paraffin-embedded sections with PSR, which stains collagen (76). Sections were deparaffinized with xylene incubations and subsequently rehydrated with ethanol incubations (100% to 70%, followed by distilled water). Sections were then incubated with PSR for 1 hour at room temperature, followed by washes in acidified water. Prior to mounting, sections were submerged in ethanol concentrations (95% to 100%), followed by xylene incubations, and then subsequently mounted with Permount mounting medium (Thermo Fisher Scientific). Sections were imaged with Brightfield microscopy on the EVOS M7000 imaging system (Thermo Fisher Scientific). Fibrotic (collagenous) regions were expressed as a percentage against standardized total regions of interest (anatomically consistent between all samples), which included whole atria and a central region of the ventricular free wall. Artefacts (such as irregular tissue folds) were avoided during the blinded analysis. Fibrosis was quantified using ImageJ (National Institute of Health).

#### Hematoxylin and Eosin (H&E)

Cardiac tissue disorganization was quantified using H&E staining, as previously described in our lab (54). Disorganized cardiac tissue was defined by loss of nuclear detail and the presence of coalescing nuclei. Sections were imaged with Brightfield microscopy on the EVOS M7000 imager. Cardiac tissue disorganization was calculated by dividing areas containing coalesced nuclei or lost nuclear detail by total region of interest (artefacts and dead space subtracted) and expressed as a percentage. Regions of interest were kept consistent anatomically between samples. Blinded analyses were completed using ImageJ.

#### Immunofluorescence

To assess inflammatory cytokine presence, and markers of fibrosis, tissue sections were incubated with primary antibodies and imaged against corresponding fluorescent secondary antibodies. Sections were deparaffinized with xylene incubations, rehydrated with gradient ethanol incubations (100% to 70%, followed by distilled water), and then incubated in boiling Tris-EDTA (TE) buffer pH 9.0 for antigen retrieval. Sections were incubated with equal parts pre-conjugated wheat germ agglutinin (WGA) (AlexaFluor 488; Cat. No. W11261; 1:1000) and multiplexed primary antibodies for IL-6 (rabbit; Cat. No. P620; 1:100) and IL-10 (mouse; Cat. No. ARC9102; 1:100) or TGF-β1 (rabbit; Cat. No. PA5-120329; 1:100) and α-SMA (mouse; Cat. No. 14-9760-82; 1:100) (Thermo Fisher Scientific), on duplicate sections per set of antibodies. The following morning, primary antibodies were rinsed off with PBS-T and sections were incubated with the following fluorescent secondary antibodies: AlexaFluor 594 (Goat Anti-Rabbit; Cat. No. ab150080; 1:1000), or Alexafluor 647 (Donkey Anti-Mouse; Cat. No. ab150107; 1:1000) (Abcam, Cambridge, UK) corresponding to the primary antibody for 1 hour at room temperature. Sections were mounted using an aqueous Fluoroshield DAPI mounting medium (Abcam) with edges sealed using clear nail polish. Slide microscopy was conducted using a fluorescence slide scanner (Zeiss AxioScan) at 20x magnification. Sections were analyzed using the QuPath-0.4.0 software. While blinded, the analyzer selectively avoided tissue folds and other fluorescent artefacts that influenced data, across all experimental groups. Entire atria were analyzed, while ROIs were kept consistent between ventricles. Endocardial, fibrotic epicardial, and septal regions were analyzed. Data was interpreted as Mean Fluorescent Intensity (MFI) corresponding to each of the four channels: AF408, AF488, AF594, and AF647, and expressed as Arbitrary Units (A.U.). The MFI from each channel of arbitrarily selected background ROIs (or neutral non-tissue sections) were averaged and subtracted from the MFI of each channel from the four chambers as a method for normalizing each section to its slide. *Minimal Feret Diameter:* To quantify cardiomyocyte size, a blinded analysis was conducted using the WGA (cellular membrane) filter on IF sections. Twenty arbitrarily selected cardiomyocyte cross-sections from the top, middle, and bottom portions of each ventricle were free hand traced, pooled together, and averaged to obtain a mean cardiomyocyte size (μm^2^) for each ventricle.

### Western blotting

Hearts were excised, promptly weighed, and subsequently flash frozen in liquid nitrogen. Heart weights were normalized to both body weight and tibial length. Western blotting was performed using standard SDS-PAGE procedures. A frozen section of each ventricle (∼10-15 mg in size for RV and ∼20-25 mg for LV) was homogenized using plastic microcentrifuge tubes and an ultrawave Sonic Dismembrator Model 500 (Thermo Fisher Scientific) in ice-cold lysis buffer containing (in mM) 20 Tris/HCl, 150 NaCl, 1 EDTA, 1 EGTA, 2.5 Na_4_O_7_P_2_, 1 Na_3_VO_4_, and 1% Triton X-100, supplemented with protease and phosphatase inhibitors (Millipore Sigma) (pH 7.0), as previously published (77,78). Protein concentrations were determined using a bicinchoninic acid (BCA) assay (Life Technologies, Carlsbad, CA, USA). Denatured and reduced protein was subjected to SDS-PAGE on 10-14% gels followed by transfer onto 0.2 μm low-fluorescence polyvinylidene difluoride membrane (Bio-Rad, Mississauga, Canada) and blocked with Li-COR Intercept Blocking Buffer for 1 hour (LI-COR, Lincoln NE, USA). Membranes were then immunoblotted overnight (4°C) with the specific primary antibodies listed below.

To quantify mitochondrial electron transport chain complex subunits (including V-ATP5A (55 kDa), III-UQCRC2 (48 kDa), IV-MTCO1 (40 kDa), II-SDHB (30 kDa), and I-NDUFB8 (20 kDa)), a commercially available monoclonal rodent OXPHOS cocktail was used (Cat. No. ab110413; 1:1000) (Abcam). Commercially available polyclonal antibodies were used to detect AMPKα (Cat. No. 2532; 62 kDa; 1:500), p-AMPKα (Thr172) (Cat. No. 2535; 62 kDa; 1:500), p38MAPK (Cat. No. 9212; 40 kDa; 1:500), p-p38MAPK (Thr180/Tyr182) (Cat. No. 9211; 43 kDa; 1:500), LC3BII/I (Cat. No. 2775S; 14/16 kDa; 1:500), and a monoclonal antibody was used to detect SQSTM1/p62 (Cat. No. 23214S; 62 kDa; 1:500) (Cell Signaling Technology, Whitby, Ontario).

Following an overnight primary antibody incubation, membranes were washed 3×5 minutes in TBS-T, and incubated at room temperature with the appropriate infrared fluorescent secondary antibody (LI-COR IRDye 680nm (Cat. No. 925-68020) or 800nm (Cat. No. 925-32214)) at previously optimized dilutions (1:10,000 to 1:20,000). Immunoreactive proteins were detected using infrared imaging (LI-COR CLx; LI-COR, Lincoln, NE, USA) and quantified on ImageJ by densitometry. All images were normalized to total protein from the same corresponding membrane using Amido Black total protein stain (Cat. No. A8181) (Millipore Sigma).

### Preparation of permeabilized muscle fibre bundles

Mitochondrial bioenergetic assays were performed using permeabilized muscle fiber bundle (PmFB) preparations from heart tissue as we have described previously (4,54,79). Upon sacrifice, the heart was immediately submerged in ice-cold ATP-containing media (BIOPS buffer) designed to mimic the extracellular environment. Tissue was carefully separated with precision forceps into small (0.6-1.6 mg) bundles that were blotted dry using a Kimwipe and weighed in 1.5 mL tared ice-cold BIOPS. Respiration values were normalized to fibre bundle wet weight. Extra precaution was taken when separating the tissue such that collagenous (non-respiring) regions were avoided when preparing bundles, however, some bundles may have had larger non-respiring regions than others. Bundles were permeabilized in BIOPS that contained saponin (40 μg/ml) for 30 mins at 4°C on a rotator, which digests the cell membrane and allows the cytosol to be washed out by selectively permeabilizing the sarcolemma and leaving other intracellular organelles intact.

### Mitochondrial respiration

High-resolution respirometry (measure of mitochondrial oxygen consumption) was performed on heart PmFBs using the Oroboros Oxygraph-2k (OROBOROS Instruments, Corp., Innsbruck, Austria). PmFBs were submerged in 2 mL Buffer Z containing (in mM) 105 K-MES, 30 KCl, 10 KH_2_PO_4_, 5 MgCl_2_ • 6H_2_O, 1 EGTA, and 5 mg/mL BSA, in the respirometer chambers in the presence of 5 μM blebbistatin to prevent contraction of PmFBs by rigor in response to ADP while stirring at 750 RPM and 37°C (79). Chambers were manually oxygenated with 100% O_2_ to an initial concentration of ∼350 μM and experiments were completed prior to chamber [O_2_] dropping past 150 μM to prevent oxygen limitations to respiration.

Oxygen consumption was measured in either Buffer Z devoid of creatine (Cr) to prevent activation of mitochondrial creatine kinase (mtCK) (creatine-independent), or in Buffer Z with 20 mM Cr to saturate mtCK and promote phosphate shuttling through this complex (creatine-dependent) as performed previously (55).

For each heart, one PmFB from each ventricle was used to assess complex-I supported respiration using pyruvate (5 mM; carbohydrate substrate) and malate (2 mM) (NADH; State II respiration) followed by titrations of basal (25 μM), sub-maximal (100 and 500 μM), and saturating ADP (1000 and 5000 μM) to assess mitochondrial responsiveness to ADP across a range of metabolic demands (State III coupled respiration) (80). Following ADP titrations, 10 mM glutamate was added to further saturate complex-I with NADH generation. Next, 10 μM cytochrome *c* was added to test mitochondrial outer membrane integrity. Anomalous cytochrome *c* responses (over 15%), where ADP-stimulated respiration was correspondingly low, were selectively removed. 10 mM succinate (FADH_2_-stimulating) was then added to support complex-II respiration (80).

In a separate PmFB from each ventricle, a fatty acid-based protocol was implemented with 10 μM L-Carnitine, 20 μM Palmitoyl-CoA, and 500 μM malate to shuttle electrons through beta-oxidation pathways (81). This protocol generates NADH (Complex I stimulation) and FADH_2_ from β-oxidation (ETF stimulation) and the TCA cycle (Complex II stimulation). A similar ADP titration protocol was then performed as described above. Following ADP titrations, 10 μM cytochrome *c* was added.

### Mitochondrial H_2_O_2_ Emission (mH_2_O_2_)

mH_2_O_2_ emission was detected spectrofluorometrically (QuantaMaster 40, HORIBA Scientific, Irvine, CA, USA) as previously described (54). Assays were conducted in Buffer Z supplemented with 10 μM Amplex Ultra Red (AUR) (an H_2_O_2_ fluorogenic probe) (Life Technologies), 1 U/mL horseradish peroxidase (HRP), 1 mM EGTA, 40 U/mL Cu-Zn SOD1, 5 μM blebbistatin, and 20 mM Cr.

Two protocols were implemented in separate PMFBs, as follows. In protocol 1, maximal mH_2_O_2_ in the absence of ADP (State II), was induced using the complex I-supporting substrates pyruvate (10 mM) and malate (2 mM) (NADH; forward electron flow). Next, stepwise ADP concentrations (25-100 μM (physiological ranges) and 500 μM (saturating for suppression of mH_2_O_2_) were titrated progressively to determine ADP’s ability to suppress mH_2_O_2_ and in the presence of 20 mM Cr (80). In protocol 2, (separate PmFB), 10 mM succinate (State II) was added to stimulate mH_2_O_2_ by reverse electron flow from complex-II (FADH_2_) to complex-I followed by ADP titrations as performed in Protocol 1. To assess the effects of creatine-dependent cycling of adenylates (ADP/ATP) on mH_2_O_2_ attenuation, the succinate protocols were repeated in a separate PmFB in the absence of creatine in the buffer given creatine enhances the ability of ADP to attenuate mH_2_O_2_ (82).

At culmination of the protocol, bundles were collected and lyophilized in a freeze-dryer (Labconco, Kansas City, MO, USA) for ∼4 hours and weighed using a microbalance (Sartorius Cubis Microbalance, Gottingen, Germany). Rate of mH_2_O_2_ was calculated using a standard curve in identical assay conditions. Values were normalized to fibre bundle dry weight.

### Calcium retention capacity

Mitochondrial calcium retention capacity (CRC) was performed as previously described (28). Prior to commencing each experiment, a cuvette lined with 10 mM EGTA in ∼500 uL deionized water was placed on a stir plate at room temperature for at least 10 mins. Water was aspirated without rinsing to allow EGTA to coat the cuvette, thus ensuring that residual Ca^2+^ in the assay buffer (Buffer Y) would be chelated. Experiments were measured spectrofluorometrically (QuantaMaster 80, HORIBA Scientific) at 37°C and continuous stirring in a cuvette with 300 μL Buffer Y containing 1 μL Calcium Green-5N (Invitrogen) as a fluorescent probe specific to Ca^2+^, 2 μM thapsigargin (to inhibit SERCA), 5 μM blebbistatin, and 40 μM EGTA. For each experiment, minimum fluorescence (Fmin) was obtained by first adding 5 mM glutamate, 2 mM malate, and 500 μM ADP to Buffer Y containing the PmFB. Following these additions, a single 8 nmol bolus of CaCl_2_ was titrated into the cuvette, followed by subsequent 4 nmol boluses until mitochondrial permeability transition (mPT) occurred, as reflected by an increase in fluorescence representing Ca^2+^ release from saturated mitochondria. Once mPT was established, two additional saturating boluses of 0.5 mM CaCl_2_ were titrated to saturate the fluorophore and establish a maximum fluorescence (Fmax). To calculate changes in free Ca^2+^ in the cuvette as an outcome of mitochondrial Ca^2+^ uptake, the known Kd for Calcium Green-5N and equations for calculating free ion concentration were utilized (54). Once an Fmax was established, PmFBs were carefully recovered and lyophilized in a freeze-dryer overnight and weighed on a microbalance for dry weight trace normalization.

### Caspase activity assay

Caspase activity was measured in whole-muscle lysates that were prepared in the absence of protease inhibitors, as described elsewhere (83). Lysis buffer consisted of (in mM): 20 HEPES, 10 NaCl, 1.5 MgCl2, 1 DTT, 20% glycerol, and 0.1% Triton X-100. Briefly, samples were incubated at room temperature in 20 μM Ac DEVD AFC (caspase 3), Ac LEHD AFC (caspase 9), or Ac-IETD-AMC (caspase-8) in assay buffer containing 20 mM HEPES, pH 7.4, 10 mM DTT, and 10% glycerol. Fluorescence was measured at room temperature using a Cytation 5 Imaging Multi-Mode Reader with excitation (400 nm) and emission (505 nm) wavelengths for AFC substrates (Caspase-3 and Caspase-9). Excitation (360 nm) and emission (440 nm) wavelengths were utilized for AMC substrates (Caspase-8). Values were normalized to protein concentration of samples and expressed as a fold change in fluorescence.

### Flow cytometry

Heparinized mice were anesthetized with isoflurane and placed into a nose cone with continuous vaporization. The RV was perfused with 7 mL pre-warmed (37°C) EDTA buffer (pH 7.8) to chelate calcium and prevent aggregation of cells. The heart was then removed and placed into a shale plate containing EDTA buffer, followed by perfusion into the LV with 10 mL EDTA to remove remaining blood. The heart was then perfused with 3.5 mL pre-warmed perfusion buffer (pH 7.8) to arrest the heart in diastole. Upon taking whole heart weight, RV and LV were surgically separated, washed with remaining perfusion buffer, and were subsequently dried and weighed. Finely minced pieces of each ventricle were digested for 60 mins at 37°C on a 140 RPM shaker in DMEM containing DNase I (60 U/mL) (Sigma; Cat no. 11284932001), collagenase type II (450 U/mL) (Worthington; Cat no. LS004176), and hyaluronidase (60 U/mL) (Sigma; Cat no. h3506). Digested chamber-specific tissue was then filtered through 100 μM cell strainers (Miltenyi Biotec) and washed with 5 mL DMEM + 20% FBS into 15 mL conical tubes. Samples then underwent centrifugation for 5 mins at 4°C and 500 RCF. With only 2 mL remaining in the conical tubes, the pellets were resuspended, and fractions were separated into ‘unstained’ and ‘full stained’ 1.5 mL tubes. Following another identical centrifugation protocol, pellets were resuspended in FACS buffer (DPBS containing 2 mL FBS stock and 1 mM EDTA). FC block (Thermo Fisher; Cat no. 14-0161-85) was added to all tubes to prevent auto-antibody binding, and tubes were incubated on a shaker at room temperature for 15 mins. Additional FACS buffer was added to all unstained tubes while equal volume cell surface marker antibody master mix was added into the full stained tubes. All tubes were kept away from light and were incubated for 30 mins at 4°C on a shaker. Samples were then centrifuged again (same protocol), resuspended in FACS buffer and intracellular (IC) fixation buffer (Thermo Fisher; Cat no. 00-8222-49), and were once again incubated for 30 mins at 4°C on a shaker. After one final centrifugation (same protocol), pellets were resuspended in 1 mL FACS buffer and stored in a dark, humidity-controlled 4°C fridge for subsequent processing using the Cytek Aurora 5-laser spectral flow cytometer (CytekBio).

### Cell surface marker antibody master mix

CD45 – 1:1200 – Thermo Fisher Cat no. 64-0451-82 – 30-F11 AF700

CD11b – 1:1000 – Thermo Fisher Cat no. 63-0112-82 – M1/70 SB600

CD64 – 1:800 - Thermo Fisher Cat no. 17-0641-82 - X54-5/7.1 APC

F4/80 – 1:600 – BioLegend Cat no. 10630 – BV650

CCR2 – 1:100 – BioLegend Cat no. 13354 - FITC

### Statistics

All results are expressed as mean ± SD. Outliers were identified and excluded in accordance with ROUT testing (Q=0.5%). Subsequently, the D’Agostino-Pearson Omnibus normality parameters were performed to determine if data resembled a Gaussian (normal), or lognormal distribution (GraphPad Prism 9.3.0 Software, La Jolla, CA, USA). Complex I-stimulated respiration in both 20 mM Cr and No Cr conditions in the LV failed normality – therefore, relevant datasets were log-transformed, and we subsequently conducted standard parametric two-way ANOVA testing. All other datasets passed normality. A two-way ANOVA was used for ADP-stimulated mitochondrial respiration and ADP-suppression of mH_2_O_2_, followed by Benjamini, Krieger, and Yekutieli’s *post hoc* analysis to correct for false discovery rate when a significant interaction was observed (84). A one-way ANOVA was conducted for morphometric, Western blot, histopathology (fibrosis and myocardial disorganization), and immunofluorescence, followed by *post hoc* analyses where relevant. Significance was established at *p* < 0.05 for all statistics. While corrected *p*-values are typically reported as q-values with the Benjamini, Krieger, and Yekutieli’s *post-hoc* analysis, we herein report them as *p*-values.

## Acknowledgements

ALY688 was provided by Allysta Pharmaceuticals. The authors would like to thank Drs. Brittany Edgett and Laura Castellani for their technical insight and assistance.

## Author contributions

Conceptualization: SG, CGRP, CAB

Methodology: SG, CGRP, PDB, MCG, CAB, ANG, BAM, ANB, SY-S, FAR, JQ, JAS, GS, AAAS, PHB, HHH

Investigation: SG, CGRP

Visualization: SG, CGRP

Supervision: SG, CGRP, JAS, GS, AAAS, PHB

Writing—original draft: SG, CGRP

Writing—review & editing: SG, CGRP, LJD, PDB, MCG, CAB, ANG, BAM, ANB, SY-S, FAR, JQ, JAS, GS, AAAS, PHB, HHH

## Data and materials availability

All data are available in the main text or the supplementary materials.

## References

1. Crisafulli S, Sultana J, Fontana A, Salvo F, Messina S, Trifirò G. Global epidemiology of Duchenne muscular dystrophy: an updated systematic review and meta-analysis. Orphanet J Rare Dis. 2020 Jun 5;15(1):141.

2. Coley WD, Bogdanik L, Vila MC, Yu Q, Van Der Meulen JH, Rayavarapu S, et al. Effect of genetic background on the dystrophic phenotype in mdx mice. Hum Mol Genet. 2016 Jan 1;25(1):130–45.

3. Putten M, Putker K, Overzier M, Adamzek WA, Pasteuning-Vuhman S, Plomp JJ, et al. Natural disease history of the D2 mdx mouse model for Duchenne muscular dystrophy. FASEB J. 2019 Jul;33(7):8110–24.

4. Hughes MC, Ramos S V., Turnbull PC, Edgett BA, Huber JS, Polidovitch N, et al. Impairments in left ventricular mitochondrial bioenergetics precede overt cardiac dysfunction and remodelling in Duchenne muscular dystrophy. J Physiol. 2020 Apr 27;598(7):1377–92.

5. Belanto JJ, Mader TL, Eckhoff MD, Strandjord DM, Banks GB, Gardner MK, et al. Microtubule binding distinguishes dystrophin from utrophin. Proc Natl Acad Sci U S A. 2014 Apr 15;111(15):5723–8.

6. Cohn RD, Campbell KP. Molecular basis of muscular dystrophies. Muscle Nerve. 2000 Oct;23(10):1456–71.

7. Cohn JN, Ferrari R, Sharpe N. Cardiac remodeling—concepts and clinical implications: a consensus paper from an international forum on cardiac remodeling. J Am Coll Cardiol. 2000 Mar;35(3):569–82.

8. Kamogawa Y. Dystrophin-deficient myocardium is vulnerable to pressure overload in vivo. Cardiovasc Res. 2001 Jun;50(3):509–15.

9. Magrath P, Maforo N, Renella P, Nelson SF, Halnon N, Ennis DB. Cardiac MRI biomarkers for Duchenne muscular dystrophy. Biomark Med. 2018 Nov 30;12(11):1271– 89.

10. Tan LB, Jalil JE, Pick R, Janicki JS, Weber KT. Cardiac myocyte necrosis induced by angiotensin II. Circ Res. 1991 Nov;69(5):1185–95.

11. Anderson KR, Sutton MGSJ, Lie JT. Histopathological types of cardiac fibrosis in myocardial disease. J Pathol. 1979 Jun 14;128(2):79–85.

12. Frankel KA, Rosser RJ. The pathology of the heart in progressive muscular dystrophy: Epimyocardial fibrosis. Hum Pathol. 1976 Jul;7(4):375–86.

13. Silva MC, Magalhães TA, Meira ZMA, Rassi CHRE, Andrade ACDS, Gutierrez PS, et al. Myocardial Fibrosis Progression in Duchenne and Becker Muscular Dystrophy: A Randomized Clinical Trial. JAMA Cardiol. 2017 Feb 1;2(2):190–9.

14. Tandon A, Villa CR, Hor KN, Jefferies JL, Gao Z, Towbin JA, et al. Myocardial Fibrosis Burden Predicts Left Ventricular Ejection Fraction and Is Associated With Age and Steroid Treatment Duration in Duchenne Muscular Dystrophy. J Am Heart Assoc. 2015 Apr 22;4(4).

15. Yuan W, Xu H, Yu L, Wen L, Xu K, Xie L, et al. Association of increased epicardial adipose tissue derived from cardiac magnetic resonance imaging with myocardial fibrosis in Duchenne muscular dystrophy: a clinical prediction model development and validation study in 283 participants. Quant Imaging Med Surg. 2024 Jan;14(1):736–48.

16. Connuck DM, Sleeper LA, Colan SD, Cox GF, Towbin JA, Lowe AM, et al. Characteristics and outcomes of cardiomyopathy in children with Duchenne or Becker muscular dystrophy: A comparative study from the Pediatric Cardiomyopathy Registry. Am Heart J. 2008 Jun;155(6):998–1005.

17. Finsterer J, Stöllberger C. The Heart in Human Dystrophinopathies. Cardiology. 2003;99(1):1–19.

18. Fayssoil A, Mansencal N, Nguyen LS, Nardi O, Yaou R Ben, Leturcq F, et al. Prognosis of Right Ventricular Systolic Dysfunction in Patients With Duchenne Muscular Dystrophy. J Am Heart Assoc. 2023 Aug 15;12(16).

19. Dual SA, Maforo NG, McElhinney DB, Prosper A, Wu HH, Maskatia S, et al. Right Ventricular Function and T1 Mapping in Boys With Duchenne Muscular Dystrophy. J Magn Reson Imaging. 2021 Nov 26;54(5):1503–13.

20. Greiner E, Breaux A, Kasten J, Seo J, Ollberding NJ, Spar D, et al. Cardiac atrial pathology in Duchenne muscular dystrophy. Muscle Nerve. 2024 May;69(5):572–9.

21. Raman S V., Cripe LH. Glucocorticoid therapy for Duchenne cardiomyopathy: A Hobson’s choice? J Am Heart Assoc. 2015 Mar 26;4(4).

22. Matthews E, Brassington R, Kuntzer T, Jichi F, Manzur AY. Corticosteroids for the treatment of Duchenne muscular dystrophy. Cochrane Database Syst Rev. 2016 May 5;2016(6).

23. Shah MNA, Yokota T. Cardiac therapies for Duchenne muscular dystrophy. Ther Adv Neurol Disord. 2023 Jan 3;16.

24. Esfahani M, Movahedian A, Baranchi M, Goodarzi MT. Adiponectin: an adipokine with protective features against metabolic syndrome. Iran J Basic Med Sci. 2015 May;18(5):430–42.

25. Sung HK, Mitchell PL, Gross S, Marette A, Sweeney G. ALY688 elicits adiponectin-mimetic signaling and improves insulin action in skeletal muscle cells. Am J Physiol Physiol. 2022 Feb 1;322(2):C151–63.

26. Abou-Samra M, Selvais CM, Boursereau R, Lecompte S, Noel L, Brichard SM. AdipoRon, a new therapeutic prospect for Duchenne muscular dystrophy. J Cachexia Sarcopenia Muscle. 2020 Apr 21;11(2):518–33.

27. Otvos L. Potential Adiponectin Receptor Response Modifier Therapeutics. Front Endocrinol (Lausanne). 2019 Aug 13;10.

28. Bellissimo CA, Gandhi S, Castellani LN, Murugathasan M, Delfinis LJ, Thuhan A, et al. The slow-release adiponectin analog ALY688-SR modifies early-stage disease development in the D2. mdx mouse model of Duchenne muscular dystrophy. Am J Physiol Physiol. 2024 Apr 1;326(4):C1011–26.

29. Li X, Zhang W, Cao Q, Wang Z, Zhao M, Xu L, et al. Mitochondrial dysfunction in fibrotic diseases. Cell Death Discov. 2020 Sep 5;6(1):80.

30. Boland BJ, Silbert PL, Groover R V., Wollan PC, Silverstein MD. Skeletal, cardiac, and smooth muscle failure in Duchenne muscular dystrophy. Pediatr Neurol. 1996 Jan;14(1):7– 12.

31. Nicholls DG, Ferguson S. Bioenergetics 4. Elsevier; 2013.

32. Steinberg GR, Kemp BE. AMPK in Health and Disease. Physiol Rev. 2009 Jul;89(3):1025–78.

33. Kulisz A, Chen N, Chandel NS, Shao Z, Schumacker PT. Mitochondrial ROS initiate phosphorylation of p38 MAP kinase during hypoxia in cardiomyocytes. Am J Physiol Cell Mol Physiol. 2002 Jun 1;282(6):L1324–9.

34. Xin X, Zhou L, Reyes CM, Liu F, Dong LQ. APPL1 mediates adiponectin-stimulated p38 MAPK activation by scaffolding the TAK1-MKK3-p38 MAPK pathway. Am J Physiol Metab. 2011 Jan;300(1):E103–10.

35. de Alvaro C, Teruel T, Hernandez R, Lorenzo M. Tumor Necrosis Factor α Produces Insulin Resistance in Skeletal Muscle by Activation of Inhibitor κB Kinase in a p38 MAPK-dependent Manner. J Biol Chem. 2004 Apr;279(17):17070–8.

36. Akimoto M, Maruyama R, Kawabata Y, Tajima Y, Takenaga K. Antidiabetic adiponectin receptor agonist AdipoRon suppresses tumour growth of pancreatic cancer by inducing RIPK1/ERK-dependent necroptosis. Cell Death Dis. 2018 Jul 23;9(8):804.

37. Wissing ER, Boyer JG, Kwong JQ, Sargent MA, Karch J, McNally EM, et al. P38α MAPK underlies muscular dystrophy and myofiber death through a Bax-dependent mechanism. Hum Mol Genet. 2014 Oct 15;23(20):5452–63.

38. Tong M, Saito T, Zhai P, Oka S-I, Mizushima W, Nakamura M, et al. Mitophagy Is Essential for Maintaining Cardiac Function During High Fat Diet-Induced Diabetic Cardiomyopathy. Circ Res. 2019 Apr 26;124(9):1360–71.

39. Bekedam FT, Goumans MJ, Bogaard HJ, de Man FS, Llucià-Valldeperas A. Molecular mechanisms and targets of right ventricular fibrosis in pulmonary hypertension. Pharmacol Ther. 2023 Apr;244:108389.

40. Rizvi F, DeFranco A, Siddiqui R, Negmadjanov U, Emelyanova L, Holmuhamedov A, et al. Chamber-specific differences in human cardiac fibroblast proliferation and responsiveness toward simvastatin. Am J Physiol Cell Physiol. 2016 Aug 1;311(2):C330–9.

41. Schlüter K-D, Kutsche HS, Hirschhäuser C, Schreckenberg R, Schulz R. Review on Chamber-Specific Differences in Right and Left Heart Reactive Oxygen Species Handling. Front Physiol. 2018 Dec 17;9.

42. Sordahl LA. Differences in mitochondrial functions from right and left ventricular myocardium of four mammalian species. Comp Biochem Physiol Part B Comp Biochem. 1976 Jan;54(3):339–42.

43. Lowes BD, Minobe W, Abraham WT, Rizeq MN, Bohlmeyer TJ, Quaife RA, et al. Changes in gene expression in the intact human heart. Downregulation of alpha-myosin heavy chain in hypertrophied, failing ventricular myocardium. J Clin Invest. 1997 Nov 1;100(9):2315–24.

44. Umar S, Lee J-H, de Lange E, Iorga A, Partow-Navid R, Bapat A, et al. Spontaneous ventricular fibrillation in right ventricular failure secondary to chronic pulmonary hypertension. Circ Arrhythm Electrophysiol. 2012 Feb;5(1):181–90.

45. Tian L, Wu D, Dasgupta A, Chen K-H, Mewburn J, Potus F, et al. Epigenetic Metabolic Reprogramming of Right Ventricular Fibroblasts in Pulmonary Arterial Hypertension. Circ Res. 2020 Jun 5;126(12):1723–45.

46. Yotsukura M, Miyagawa M, Tsua T, Ishihara T, Ishikawa K. Pulmonary hypertension in progressive muscular dystrophy of the Duchenne type. Jpn Circ J. 1988;52(4):321–6.

47. Sisson TM, Sublett-Smith J, Dupont E, Hirsch R, Lorts A, Villa C. Wireless Pulmonary Artery Pressure Monitor Implantation in a Patient with Duchenne Muscular Dystrophy. Pediatr Cardiol. 2024 Jun 19;45(5):1151–3.

48. Meyers TA, Townsend D. Early right ventricular fibrosis and reduction in biventricular cardiac reserve in the dystrophin-deficient mdx heart. Am J Physiol Circ Physiol. 2015 Feb 15;308(4):H303–15.

49. Barbin ICC, Pereira JA, Bersan Rovere M, de Oliveira Moreira D, Marques MJ, Santo Neto H. Diaphragm degeneration and cardiac structure in mdx mouse: potential clinical implications for Duchenne muscular dystrophy. J Anat. 2016 May 29;228(5):784–91.

50. Thrush PT, Allen HD, Viollet L, Mendell JR. Re-examination of the Electrocardiogram in Boys With Duchenne Muscular Dystrophy and Correlation With Its Dilated Cardiomyopathy. Am J Cardiol. 2009 Jan;103(2):262–5.

51. De Giorgio D, Novelli D, Motta F, Cerrato M, Olivari D, Salama A, et al. Characterization of the Cardiac Structure and Function of Conscious D2.B10-Dmdmdx/J (D2-mdx) mice from 16-17 to 24-25 Weeks of Age. Int J Mol Sci. 2023 Jul 22;24(14):11805.

52. Tang L, Shao S, Wang C. Electrocardiographic features of children with Duchenne muscular dystrophy. Orphanet J Rare Dis. 2022 Aug 20;17(1):320.

53. Shah AM, Jefferies JL, Rossano JW, Decker JA, Cannon BC, Kim JJ. Electrocardiographic abnormalities and arrhythmias are strongly associated with the development of cardiomyopathy in muscular dystrophy. Hear Rhythm. 2010 Oct;7(10):1484–8.

54. Hughes MC, Ramos S V., Turnbull PC, Rebalka IA, Cao A, Monaco CMF, et al. Early myopathy in Duchenne muscular dystrophy is associated with elevated mitochondrial H 2 O 2 emission during impaired oxidative phosphorylation. J Cachexia Sarcopenia Muscle. 2019 Jun 2;10(3):643–61.

55. Bellissimo CA, Delfinis LJ, Hughes MC, Turnbull PC, Gandhi S, DiBenedetto SN, et al. Mitochondrial creatine sensitivity is lost in the D2. mdx model of Duchenne muscular dystrophy and rescued by the mitochondrial-enhancing compound Olesoxime. Am J Physiol Physiol. 2023 May 1;324(5):C1141–57.

56. Heywood JT, Seethala S, Khan T, Johnson A, Smith M, Rubenson D, et al. Left atrial diastolic dysfunction and pulmonary venous hypertension in atrial fibrillation: Clinical, hemodynamic and echocardiographic characteristics. J Atr Fibrillation. 2014;7(3):1117.

57. Hakimi M, Burnham T, Ramsay J, Cheung JW, Goyal NA, Jefferies JL, et al. Electrophysiologic and cardiovascular manifestations of Duchenne and Becker muscular dystrophies. Hear Rhythm. 2024;24(02882–0):S1547–5271.

58. Huang Z, Sung HK, Yan X, He S, Jin L, Wang Q, et al. The adiponectin-derived peptide ALY688 protects against the development of metabolic dysfunction-associated steatohepatitis. Clin Transl Sci. 2024;6(e13760).

59. Lone AH, Tang J, Pignalosa A, Hsu HH, Sater AAA, Sweeney G. A novel blood-based bioassay to monitor adiponectin signaling. Int Immunopharmacol. 2024;132(111890).

60. Gandhi S, Sweeney G, Perry CGR. Recent Advances in Pre-Clinical Development of Adiponectin Receptor Agonist Therapies for Duchenne Muscular Dystrophy. Biomedicines. 2024;12(7):1407.

61. Mojumdar K, Liang F, Giordano C, Lemaire C, Danialou G, Okazaki T, et al. Inflammatory monocytes promote progression of Duchenne muscular dystrophy and can be therapeutically targeted via CCR 2. EMBO Mol Med. 2014 Nov 13;6(11):1476–92.

62. Gabbiani G. The role of contractile proteins in wound healing and fibrocontractive diseases. Methods Achiev Exp Pathol. 1979;9:187–206.

63. Hinz B. Myofibroblasts. Exp Eye Res. 2016 Jan;142:56–70.

64. Tomasek JJ, Gabbiani G, Hinz B, Chaponnier C, Brown RA. Myofibroblasts and mechano-regulation of connective tissue remodelling. Nat Rev Mol Cell Biol. 2002 May 1;3(5):349– 63.

65. Shinde A V., Humeres C, Frangogiannis NG. The role of α-smooth muscle actin in fibroblast-mediated matrix contraction and remodeling. Biochim Biophys Acta - Mol Basis Dis. 2017 Jan;1863(1):298–309.

66. Nitahara-Kasahara Y, Hayashita-Kinoh H, Chiyo T, Nishiyama A, Okada H, Takeda S, et al. Dystrophic mdx mice develop severe cardiac and respiratory dysfunction following genetic ablation of the anti-inflammatory cytokine IL-10. Hum Mol Genet. 2014 Aug 1;23(15):3990–4000.

67. Frangogiannis NG. Transforming growth factor–β in tissue fibrosis. J Exp Med. 2020 Mar 2;217(3).

68. Gandhi S, Sweeney HL, Hart C, Han R, Perry CGR. Cardiomyopathy in Duchenne Muscular Dystrophy and the Potential for Mitochondrial Therapeutics to Improve Treatment Response. Cells. 2024;13(14):1168.

69. Bellissimo CA, Garibotti MC, Perry CGR. Mitochondrial stress responses in Duchenne muscular dystrophy: metabolic dysfunction or adaptive reprogramming? Am J Physiol Physiol. 2022 Sep 1;323(3):C718–30.

70. Bernardi P, Gerle C, Halestrap AP, Jonas EA, Karch J, Mnatsakanyan N, et al. Identity, structure, and function of the mitochondrial permeability transition pore: controversies, consensus, recent advances, and future directions. Cell Death Differ. 2023 Aug 17;30(8):1869–85.

71. Araya LE, Soni I V., Hardy JA, Julien O. Deorphanizing Caspase-3 and Caspase-9 Substrates In and Out of Apoptosis with Deep Substrate Profiling. ACS Chem Biol. 2021 Nov 19;16(11):2280–96.

72. Kuno A, Hosoda R, Sebori R, Hayashi T, Sakuragi H, Tanabe M, et al. Resveratrol Ameliorates Mitophagy Disturbance and Improves Cardiac Pathophysiology of Dystrophin-deficient mdx Mice. Sci Rep. 2018 Oct 22;8(1):15555.

73. Kang C, Badr MA, Kyrychenko V, Eskelinen EL, Shirokova N. Deficit in PINK1/PARKIN-mediated mitochondrial autophagy at late stages of dystrophic cardiomyopathy. Cardiovasc Res. 2018;114(1):90–102.

74. Mao X, Kikani CK, Riojas RA, Langlais P, Wang L, Ramos FJ, et al. APPL1 binds to adiponectin receptors and mediates adiponectin signalling and function. Nat Cell Biol. 2006 May 1;8(5):516–23.

75. Yamauchi T, Kamon J, Ito Y, Tsuchida A, Yokomizo T, Kita S, et al. Cloning of adiponectin receptors that mediate antidiabetic metabolic effects. Nature. 2003 Jun 12;423(6941):762–9.

76. Junqueira LCU, Bignolas G, Brentani RR. Picrosirius staining plus polarization microscopy, a specific method for collagen detection in tissue sections. Histochem J. 1979 Jul;11(4):447–55.

77. Houde VP, Donzelli S, Sacconi A, Galic S, Hammill JA, Bramson JL, et al. AMPK β1 reduces tumor progression and improves survival in p53 null mice. Mol Oncol. 2017 Sep 28;11(9):1143–55.

78. Delfinis LJ, Bellissimo CA, Gandhi S, DiBenedetto SN, Garibotti MC, Thuhan AK, et al. Muscle weakness precedes atrophy during cancer cachexia and is linked to muscle-specific mitochondrial stress. JCI Insight. 2022 Dec 22;7(24).

79. Perry CGR, Kane DA, Lin C-T, Kozy R, Cathey BL, Lark DS, et al. Inhibiting myosin-ATPase reveals a dynamic range of mitochondrial respiratory control in skeletal muscle. Biochem J. 2011 Jul 15;437(2):215–22.

80. Perry CGR, Heigenhauser GJF, Bonen A, Spriet LL. High-intensity aerobic interval training increases fat and carbohydrate metabolic capacities in human skeletal muscle. Appl Physiol Nutr Metab. 2008 Dec;33(6):1112–23.

81. Cooper MA, McCoin C, Pei D, Thyfault JP, Koestler D, Wright DE. Reduced mitochondrial reactive oxygen species production in peripheral nerves of mice fed a ketogenic diet. Exp Physiol. 2018 Sep 8;103(9):1206–12.

82. Meyer LE, Machado LB, Santiago APSA, Da-Silva WS, De Felice FG, Holub O, et al. Mitochondrial Creatine Kinase Activity Prevents Reactive Oxygen Species Generation. J Biol Chem. 2006 Dec;281(49):37361–71.

83. Rahman FA, Hian Cheong DJ, Boonstra K, Ma A, Thoms JP, Zago AS, et al. Augmented mitochondrial apoptotic signaling impairs C2C12 myoblast differentiation following cellular aging through sequential passaging. J Cell Physiol. 2024 Jan 11;

84. Glickman ME, Rao SR, Schultz MR. False discovery rate control is a recommended alternative to Bonferroni-type adjustments in health studies. J Clin Epidemiol. 2014 Aug;67(8):850–7.

