## Supplementary material for "Adiponectin-receptor agonism prevents right ventricular tissue pathology in a mouse model of Duchenne muscular dystrophy": Figure S1

Shivam Gandhi *et al.*

**This PDF file includes:**

Supplementary Figures

Figs. S1 to S8

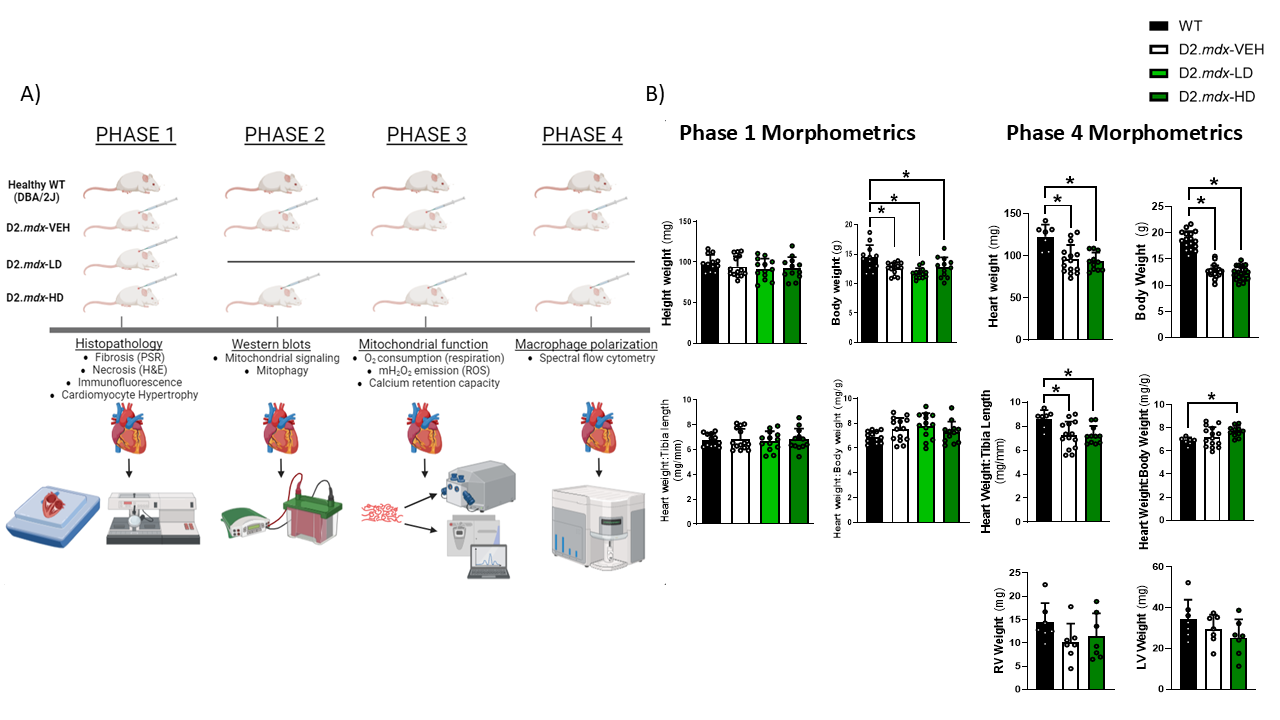

**Figure S1. Study design and corresponding timeline of data collection phases, with morphometric data from 4-week-old WT, D2.*mdx-*LD (phase 1 only), D2.*mdx*-VEH, and D2.*mdx*-HD.** A) To collect chamber-specific data for all measures with a sufficient sample size, four phases of in-house breeding and daily injections were required. D2.*mdx*-LD mice were dropped following the first phase due to logistical considerations. B) Specific comparisons include heart weight (mg), body weight (g), heart weight normalized to tibia length (mg/mm), heart weight normalized to body weight (mg/g), and ventricle-specific weights (phase 4 only). Results represent mean ± SD; n=7-14. All *p* values are FDR-adjusted by Benjamini, Krieger, and Yekutieli *post-hoc* analyses. **p*<0.05 denotes significance.

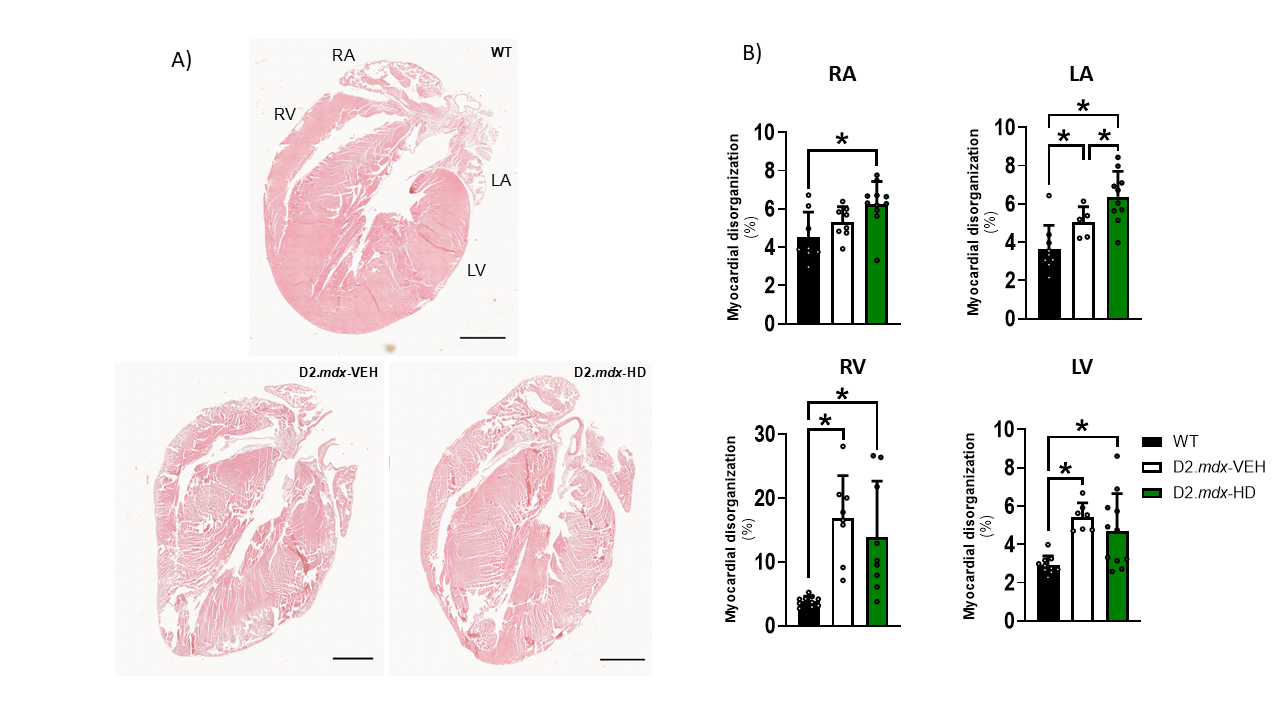

**Figure S2. D2*.mdx* mice demonstrate robust myocardial disorganization across various chambers which is not rescued by ALY688.** A) H&E staining on 5 μM thick frontal sections of paraffin-embedded cardiac tissue reveals myocardial disorganization (represented as a % of total tissue area analyzed), which is defined by loss of cellular/nuclear detail and the presence of coalescing nuclei. B) Quantitative analysis of four-chamber myocardial disorganization. Comparable results were demonstrated within the LV in D2.*mdx*-VEH. Images were taken with EVOS M7000 Imaging System at 20x magnification. Results represent mean ± SD; n=5-11. Scale bars are 1 mm. All *p* values are FDR-adjusted by Benjamini, Krieger, and Yekutieli *post-hoc* analyses. **p*<0.05 denotes significance.

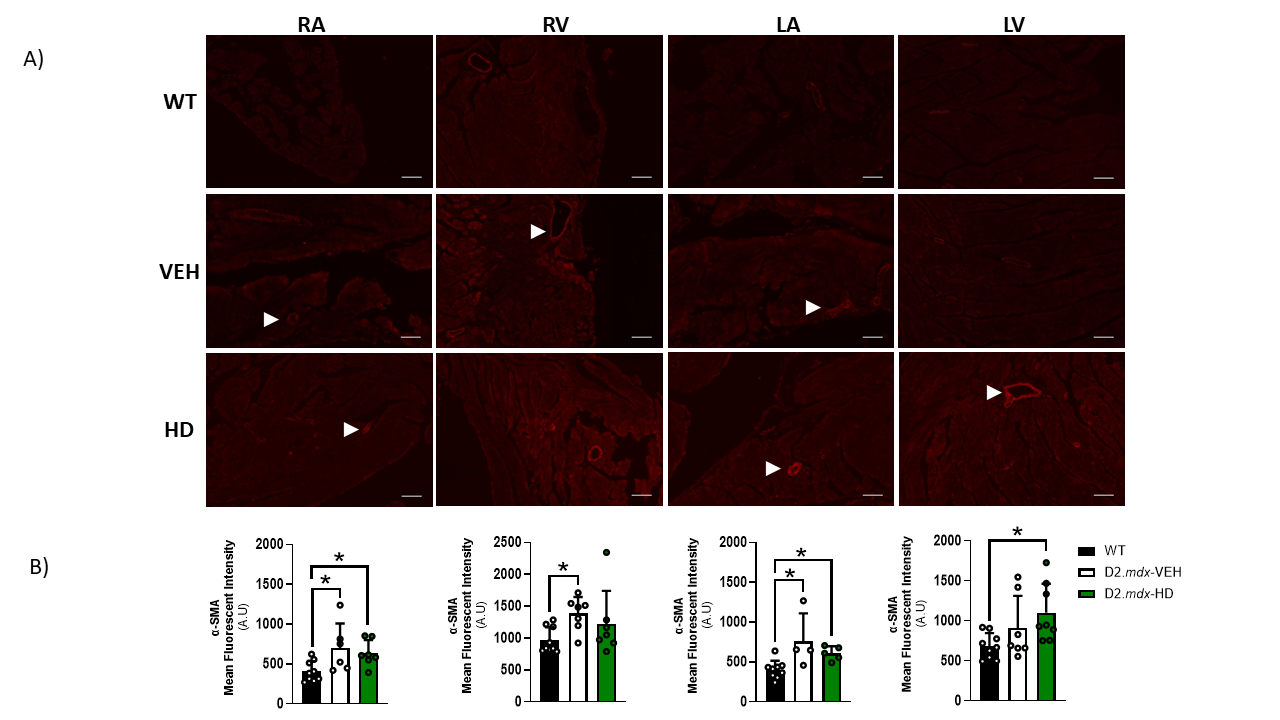
**Figure S3. Chamber-specific immunofluorescent staining reveals distinct differences in α-SMA across groups in D2*.mdx* mice.** A) α-SMA (denoted by white arrows) was quantified by Mean Fluorescent Intensity (A.U.) per region of interest (B). Whole atria and free-wall segments of the ventricles were analyzed. Stains were conducted on 5 μm thick paraffin-embedded frontal sections and imaged by confocal microscopy at 20x magnification. Results represent mean ± SD; n=4-9. Scale bars are 100 μm. All *p* values are FDR-adjusted by Benjamini, Krieger, and Yekutieli *post-hoc* analyses. **p*<0.05 denotes significance. WT = Wildtype; VEH = vehicle (saline)-treated *mdx*; HD = high dose (ALY688)-treated *mdx*; RA = right atrium; LA = left atrium; RV = right ventricle; LV = left ventricle.

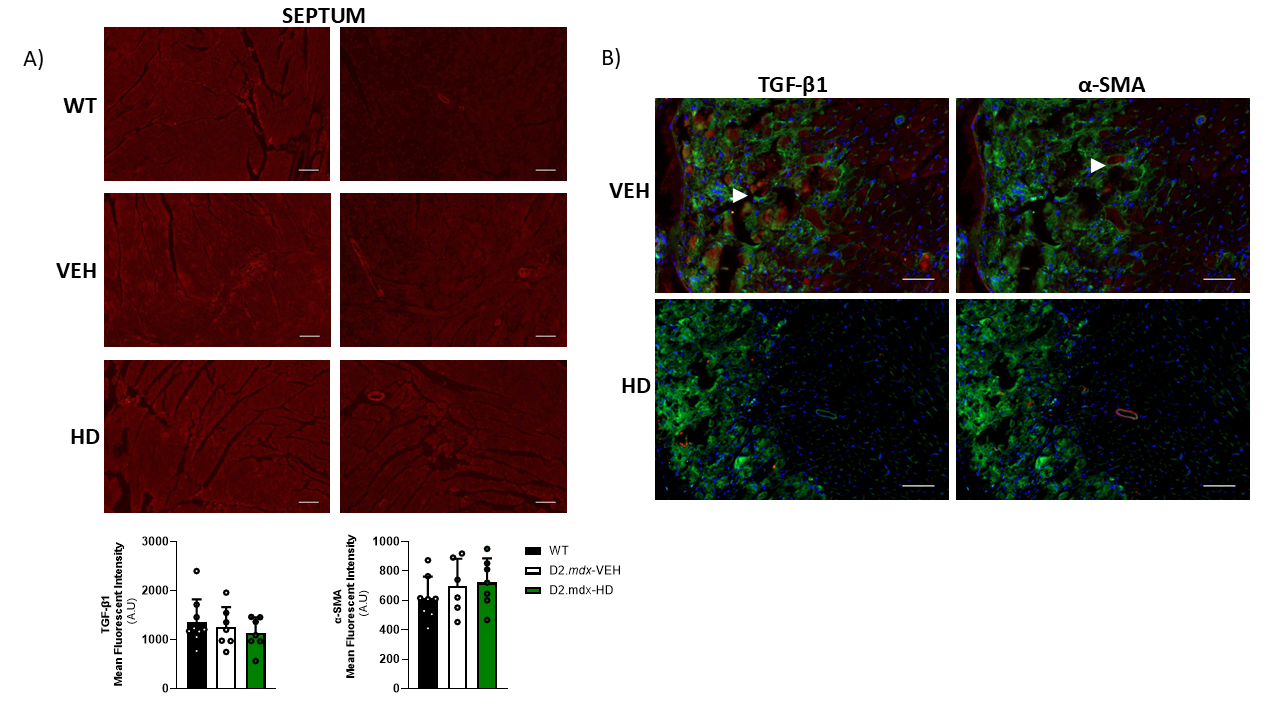

**Figure S4. Interventricular septum-specific and RV fibrotic region immunofluorescent staining of TGF-β1 and α-SMA.** A) No differences were detected in TGF-β1 or α-SMA when assessing the interventricular septum between groups. B) Presence of TGF-β1 or α-SMA (denoted by white arrows) in the fibrotic (epicardial RV) regions of D2.*mdx*-VEH and D2.*mdx*-HD mice. Statistics could not be assessed (B) due to low sample size of D2.*mdx*-HD with fibrosis. N=6-10 (septum). Scale bars are 100 μm. WT = Wildtype; VEH = vehicle (saline)-treated *mdx*; HD = high dose (ALY688)-treated *mdx*.

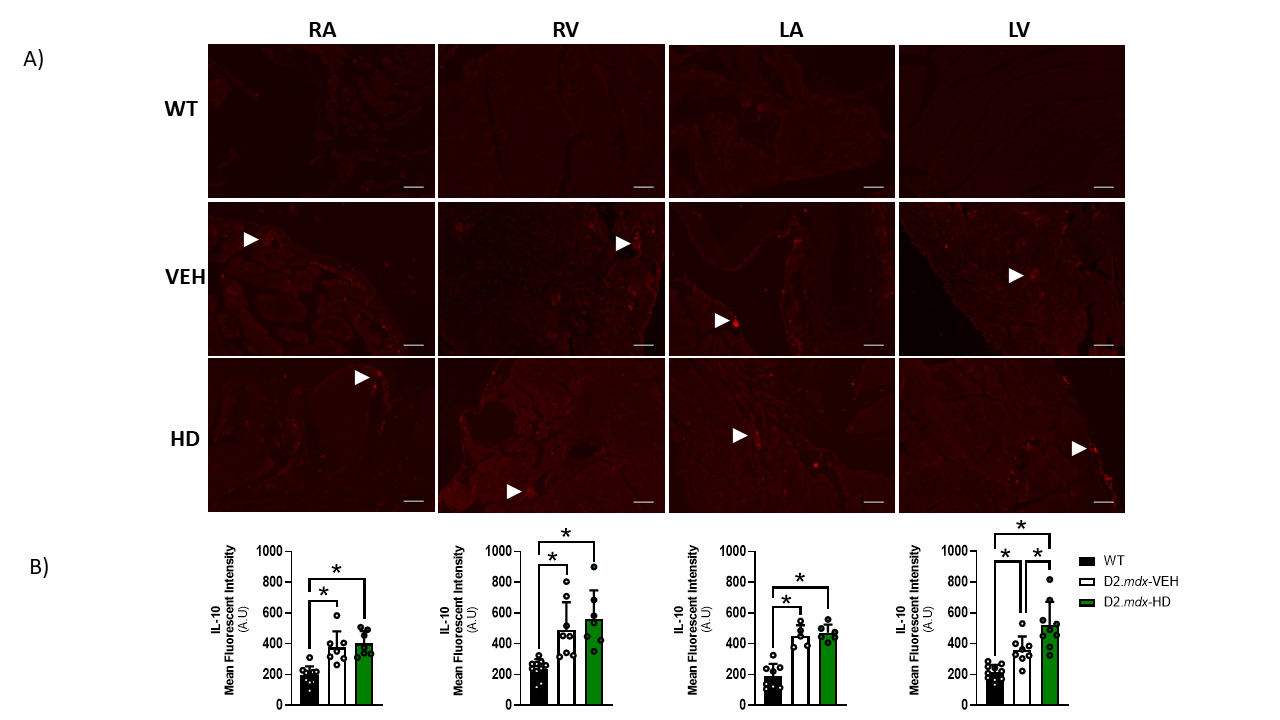

**Figure S5. Chamber-specific immunofluorescent staining of IL-10*.*** A) IL-10 (denoted by white arrows) was quantified by Mean Fluorescent Intensity (A.U.) per region of interest (B). Whole atria and free-wall segments of the ventricles were analyzed. Stains were conducted on 5 μm thick paraffin-embedded frontal sections and imaged by confocal microscopy at 20x magnification. Results represent mean ± SD; n=5-10. Scale bars are 100 μm. All *p* values are FDR-adjusted by Benjamini, Krieger, and Yekutieli *post-hoc* analyses. **p*<0.05 denotes significance. WT = Wildtype; VEH = vehicle (saline)-treated *mdx*; HD = high dose (ALY688)-treated *mdx*; RA = right atrium; LA = left atrium; RV = right ventricle; LV = left ventricle.

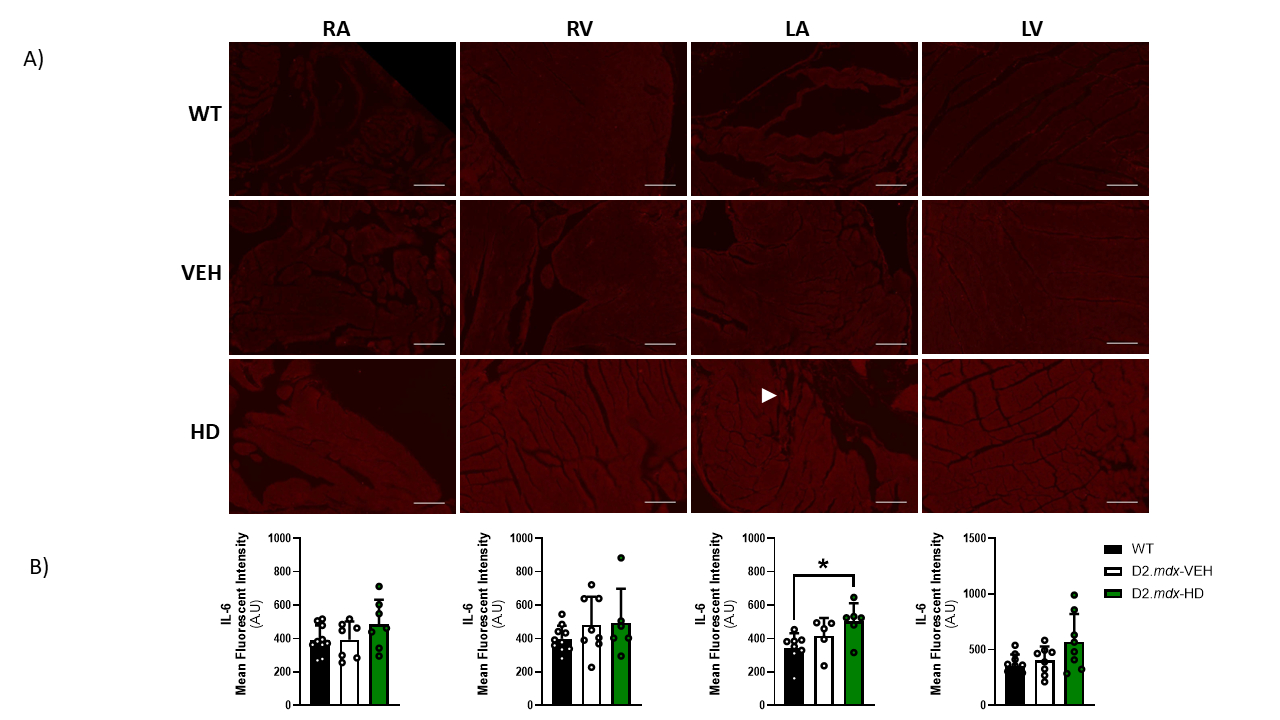

**Figure S6. Chamber-specific immunofluorescent staining of IL-6.** A) IL-6 (denoted by white arrow) was quantified by Mean Fluorescent Intensity (A.U.) per region of interest (B). Whole atria and free-wall segments of the ventricles were analyzed. Stains were conducted on 5 μm thick paraffin-embedded frontal sections and imaged by confocal microscopy at 20x magnification. Results represent mean ± SD; n=5-10. Scale bars are 100 μm. All *p* values are FDR-adjusted by Benjamini, Krieger, and Yekutieli *post-hoc* analyses. **p*<0.05 denotes significance. WT = Wildtype; VEH = vehicle (saline)-treated *mdx*; HD = high dose (ALY688)-treated *mdx*; RA = right atrium; LA = left atrium; RV = right ventricle; LV = left ventricle.

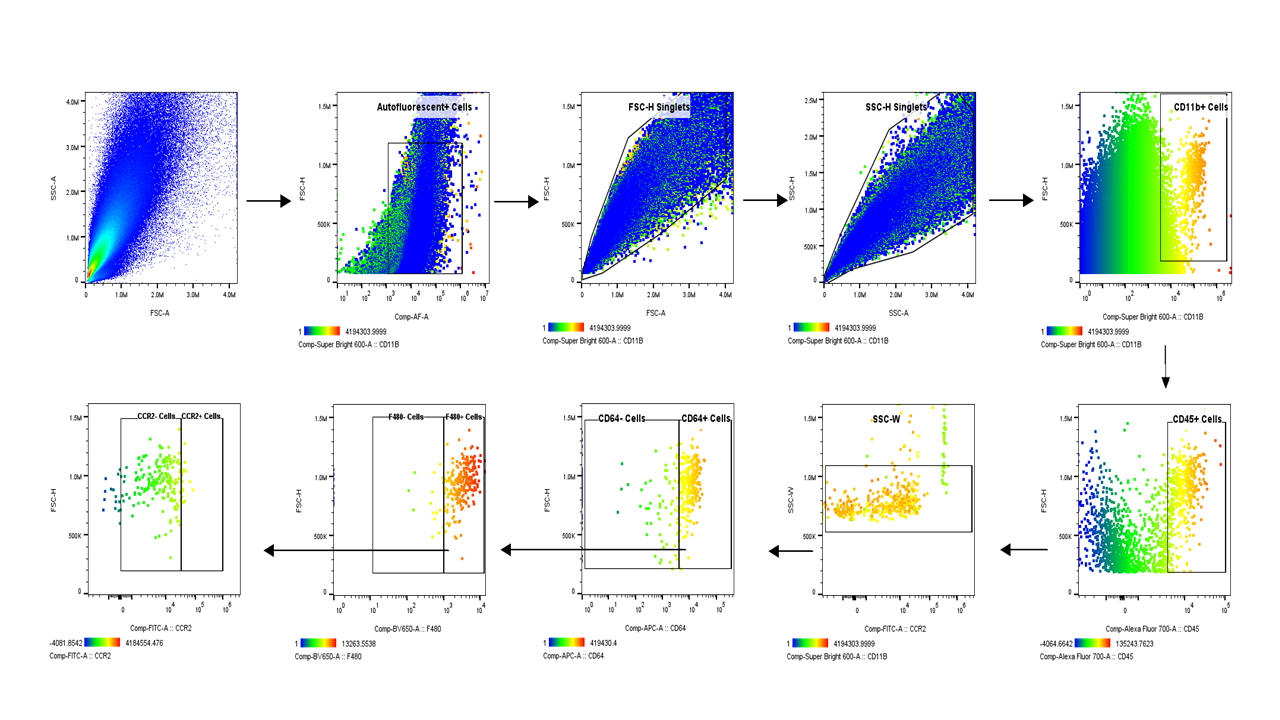

**Figure S7. Representative spectral flow cytometry gating scheme demonstrating how specific immune cell populations were gated.** Gating scheme includes several quality control checks to exclude cell aggregates and debris which may contribute to artefacts in cell count. The corresponding serotype for this gating scheme is CD11b^+^CD45^+^ CD64^+/-^F4/80^+/-^CCR2^+/-^, retrieved from a D2.*mdx*-HD RV.

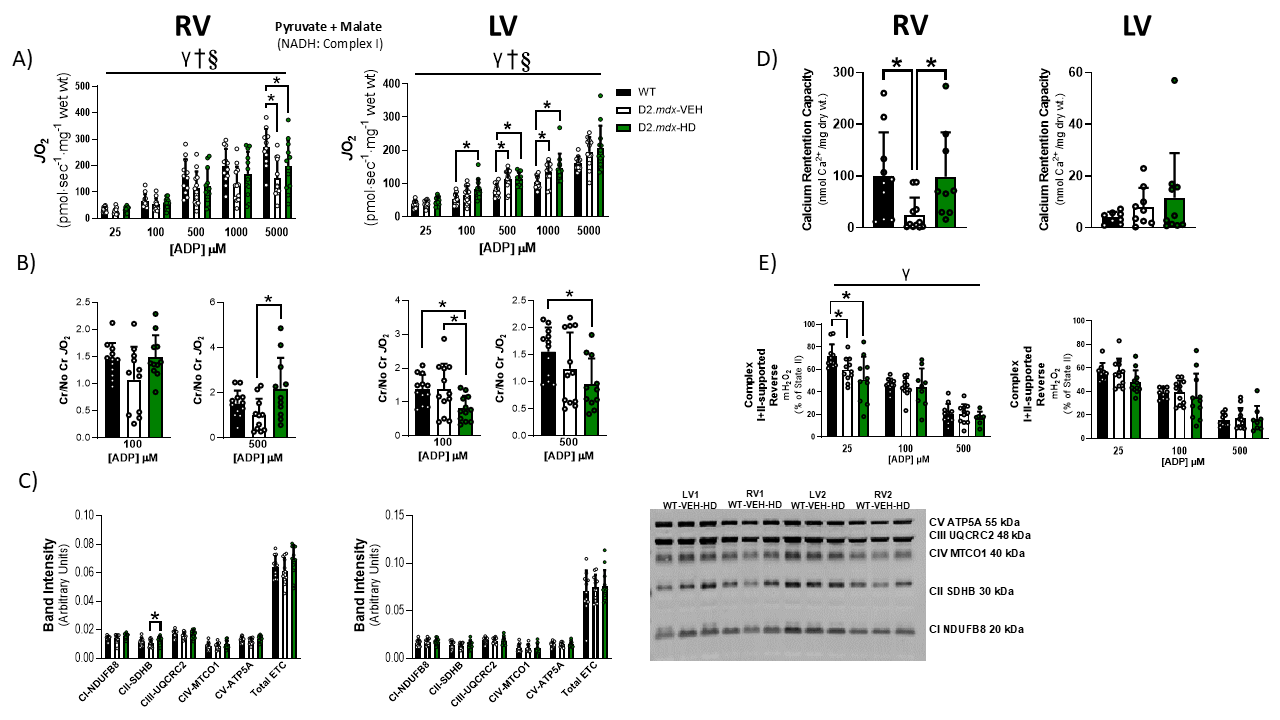

**Figure S8. Mitochondrial respiratory control by ADP during oxidative phosphorylation is protected by ALY688 in the absence of creatine, while select mitochondrial stress responses (CRC and H_2_O_2_) differ between ventricles and are only partially protected by ALY688.** A) ADP-stimulated respiration was supported by pyruvate (5 mM) supplemented with malate (2 mM) (NADH, complex-I stimulation) as an index of carbohydrate oxidation in the absence of creatine. Results represent mean ± SD; n=10-12. LV data was log-transformed (for statistical analyses) given that it did not pass normality testing. **p*<0.05 denotes significance. Main effects are denoted by horizontal bar - γ*p*<0.05 WT vs D2*.mdx*-VEH*;* §*p*<0.05 D2.*mdx*-VEH vs D2.*mdx*-HD; †*p*<0.05 WT vs D2*.mdx*-HD. B) Ventricle-specific creatine sensitivity using 100 and 500 μM ADP reveals a complex relationship between groups. n=10-12. **p*<0.05 denotes significance. C) OXPHOS content Western blot of ETC subunits in RV and LV. N=11-12. Results represent mean ± SD. **p*<0.05 denotes significance. D) Mitochondrial calcium retention capacity as an index of the potential for triggering permeability transition was assessed in permeabilized muscle fibre bundles from RV and LV. Results represent mean ± SD; n=8-11. **p*<0.05 denotes significance. E) Analysis of mitochondrial complex I+II-supported (succinate) reverse electron transfer H_2_O_2_ emission (represented as % of State II) in the presence of creatine in both ventricles across metabolic demands (25, 100, 500 μM ADP). Results represent mean ± SD; n=8-12. **p*<0.05 denotes significance; main effects are denoted by horizontal bar: γ*p*<0.05 WT vs D2*.mdx*-VEH. All *p* values are FDR-adjusted by Benjamini, Krieger, and Yekutieli *post-hoc* analyses.
